# A deep-learning-based score to evaluate multiple sequence alignments

**DOI:** 10.64898/2026.02.02.703429

**Authors:** Nimrod Serok, Ksenia Polonsky, Haim Ashkenazy, Itay Mayrose, Jeffrey L. Thorne, Tal Pupko

**Author notes:** These two authors equally contributed to this work (Nimrod Serok, Ksenia Polonsky).

## Abstract

Multiple sequence alignment (MSA) inference is a central task in molecular evolution and comparative genomics, and the reliability of downstream analyses, including phylogenetic inference, depends critically on alignment quality. Despite this importance, most widely used MSA methods optimize the sum-of-pairs (SP) score, and relatively little attention has been paid to whether this objective function accurately reflects alignment accuracy. Here, we evaluate the performance of the SP score using simulated and empirical benchmark alignments. For each dataset, we compare alternative MSAs derived from the same unaligned sequences and quantify the relationship between their SP scores and their distances from a reference alignment. We show that the alignment with the optimal SP score often does not correspond to the most accurate alignment. To address this limitation, we develop deep-learning-based scoring functions that integrate a collection of MSA features. We first introduce *Model 1*, a regression model that predicts the distance of a given MSA from the reference alignment. Across simulated and empirical datasets, this learned score correlates more strongly with true alignment accuracy than the SP score. However, *Model 1* is less effective at identifying the best alignment among alternatives. We therefore develop *Model 2*, which takes as input a set of alternative MSAs generated from the same sequences and predicts their relative ranking. *Model 2* more accurately identifies the top-ranking MSA than the SP score, *Model 1*, and several widely used alignment programs. Using simulations, we show that selecting MSAs based on our approach leads to more accurate phylogenetic reconstructions.

## Introduction

Sequence alignment involves inferring sequence homology at the residue level, meaning that when two residues are aligned, they hypothetically evolved from a common ancestral residue. Reliable alignment is therefore essential for accurately modeling evolutionary processes such as substitutions and indels (Redelings et al. 2024). As a result, sequence alignments are imperative to a wide range of bioinformatics and evolutionary genomics analyses, including phylogenetic inference, molecular dating, ancestral sequence reconstruction, detection of selective forces, and identification of remote homology (Kemena and Notredame 2009; Thompson et al. 2011). They also play a key role in structural biology—for example, in AI-based protein structure prediction tools, such as AlphaFold (Jumper et al. 2021). Beyond individual proteins or genes, sequence alignments are also used to align entire genomes and investigate large-scale evolutionary events, such as lateral gene transfer (Roettger et al. 2009). Additionally, they play a crucial role in studying the genetic basis of various diseases, including Coronavirus disease (COVID-19) (Pekar et al. 2022), speech and language disorders (Enard et al. 2002), and cancer (Tollis et al. 2017; Gassner et al. 2018).

For aligning pairs of sequences (pairwise alignment), dynamic programming algorithms are commonly used to efficiently explore the space of all possible alignments and identify those that maximize a predefined alignment score (Needleman and Wunsch 1970; Sankoff 1972). A widely adopted scoring scheme is the affine gap penalty model, which assigns distinct costs for substitutions, gap openings, and gap extensions (Gotoh 1982). For protein sequences, substitution scores are typically drawn from empirical matrices, such as PAM and BLOSUM (reviewed in Trivedi and Nagarajaram 2020). Classic dynamic programming algorithms are relatively efficient, requiring computational effort proportional to the product of the lengths of the two sequences being aligned. However, with the rapid growth of sequence databases and the increasing demand for fast and reliable alignment of query sequences against these datasets, substantial effort has gone into developing high-speed heuristic algorithms for pairwise alignment, such as BLAST (Altschul et al. 1990), MMseqs2 (Mirdita et al. 2019), and DIAMOND (Buchfink et al. 2015). Additionally, statistical alignment algorithms that incorporate explicit models of sequence evolution have been developed (Thorne et al. 1991; Hein et al. 2000). These methods generally offer higher accuracy compared to traditional dynamic programming but are computationally intensive, often several orders of magnitude slower, and are thus rarely used.

The generalization of pairwise alignment algorithms to multiple sequence alignment (MSA) necessitated updating the scoring function to accommodate multiple sequences. The most widely used scoring function for MSAs is the ‘sum-of-pairs’ (SoP) score (Carrillo and Lipman 1988), which is simply the sum of the induced pairwise alignment scores over all pairs of sequences. Specifically, consider an MSA with *N* sequences, and let *S_ij_* be the score of the induced pairwise alignment between sequences *i*, and *j*. Then 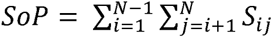.

The dynamic programming algorithm for pairwise alignment can be extended to MSAs, optimizing the SoP score. However, its computational time increases exponentially with the number of sequences (Wang and Jiang 1994). These extensive computational demands have led to a large number of heuristic algorithms that implement various optimization strategies. Most common aligners, such as Clustal Omega (Sievers and Higgins 2021), MAFFT (Katoh and Standley 2013), and PRANK (Löytynoja 2021), include a progressive alignment component, in which the MSA is iteratively built by uniting sequences according to a guide tree (Feng and Doolittle 1987). The guide tree is reconstructed without relying on the MSA, e.g., by computing pairwise distances for each pair of sequences and applying distance-based tree reconstruction methods such as the neighbor-joining (Saitou and Nei 1987). Additional heuristic strategies applied to the problem of optimizing the SoP score include iterative refinement (Hirosawa et al. 1995), divide-and-conquer (Zaharias et al. 2023), simulated annealing (Kim et al. 1994), and genetic algorithms (Gondro and Kinghorn 2007).

While many studies have focused on optimizing the SoP score, relatively few have sought to replace it with more predictive alternatives. The SoP score considers only pairwise relationships between sequences, ignoring higher-order features that quantify the conservation of entire columns. The column score, which is the percentage of columns without any substitutions, is one such feature. The information-theoretical entropy of a column, defined as ∑*p_i_log*_2_*(p_i_*), in which *p_i_* is the frequency of amino acid *i* in that column, captures more nuance conservation patterns at each column. The sum of the column-entropy scores over all alignment columns, denoted as the entropy score (ES), provides a global conservation across the entire MSA (Shenkin et al. 1991). Similarly, Hertz and Stormo (1999) proposed maximizing the information content of an MSA, defined as the extent to which observed residue frequencies in each column deviate from those expected under a background distribution, such as the overall character frequencies in the MSA. Likewise, norMD quantifies column conservation as an alternative MSA objective function (Thompson et al. 2001). Building on this concept, Nguyen and Pan (2011) introduced a novel column-scoring method that demonstrated improved performance over previous approaches.

An alternative set of scores aimed to integrate biological understanding, including structural and functional characteristics of the sequences being aligned, e.g., accounting for contact residues within the 3D structure (Kemena et al. 2011) or proportion of amino acids located in transmembrane regions and functional annotations of the aligned proteins (Ortuño et al. 2015). Orobitg et al. (2013) developed an objective function that combines several previously suggested scores, e.g., SoP scores with different gap penalties and the Triplet score used by the T-Coffee aligner (Notredame et al. 2000) that measures the consistency between the MSA and a reference library of pairwise alignments. Finally, the ClustalW algorithm (Thompson et al. 1994) modifies the SoP score to better reflect biological context by reducing the gap penalties in regions enriched with hydrophilic amino acids, which are often located on protein surfaces, where indels are more frequent.

A critical challenge in developing novel MSA scoring functions lies in the methodology used to evaluate and compare competing functions. Effective benchmarking requires two key components: (1) a set of reference MSAs, and (2) a metric that quantifies the level of disagreement between a predicted MSA and the reference alignment. However, establishing reliable benchmarks for assessing the performance of alignment methods is inherently difficult (Warnow 2021), and these challenges similarly affect the evaluation of MSA scoring functions. Two main strategies are used for generating benchmark MSAs: (1) simulation-based datasets and (2) empirical datasets derived from structural alignments. Each approach has its own strengths and limitations. Simulation-based benchmarking offers several advantages. Simulated MSAs can be produced in large quantities, the reference alignment is known, and a diverse range of evolutionary scenarios can be modeled (Iantorno et al. 2014). To ensure biological realism, simulations are typically parameterized using empirical data—for example, sequences are evolved along phylogenetic trees inferred from real datasets, with branch lengths, rate variation across sites, and indel processes modeled to reflect observed biological patterns. However, a significant limitation of simulation-based benchmarks is their reliance on simplified evolutionary models that may not accurately capture important aspects of molecular evolution. For instance, most simulations assume site independence given a phylogenetic tree, whereas in reality, protein structure and function often impose dependencies among sites.

The second benchmarking strategy involves the use of empirical protein alignments informed by structural data. While these datasets aim to reflect biologically meaningful alignments, they are still the result of inference and may not represent the true evolutionary history. Indeed, studies have shown that different structural alignment algorithms can yield inconsistent results (Hasegawa and Holm 2009; Sierk et al. 2010). Edgar (2010) noted that “questionable alignments are found in all benchmarks, especially in BAliBASE,” one of the most widely used reference databases (Thompson et al. 2005). Furthermore, structural similarity can arise through convergent evolution, leading to alignments of structurally similar—but not homologous—regions, which undermines the evolutionary validity of the benchmark (Iantorno et al. 2014). Structural benchmarks are also limited in scope: they primarily include proteins with well-defined, conserved structural regions, excluding unstructured protein regions and non-coding DNA alignments. In addition, these datasets lack scalability, particularly for applications such as machine learning, which require large and diverse training sets. Given the distinct advantages and limitations of both simulation-based and structure-based benchmarking strategies, it is advisable to evaluate alignment algorithms, including scoring functions, using both approaches.

A key component of benchmarking MSA methods is a metric that quantifies the disagreement between a predicted alignment and a reference (“reference”) alignment. The most commonly used metrics are the Total Column Similarity (TCS) score and the Residue-Pairs Similarity (RPS) score (Thompson et al. 1999). The TCS score measures the proportion of correctly recovered columns from the reference alignment, while the RPS score quantifies the proportion of correctly aligned residue pairs between the predicted and reference alignments (Thompson et al. 2005). Notably, neither the 1-TCS nor 1-RPS score is a true distance metric, as both fail to satisfy key mathematical properties of distance functions, such as symmetry and the triangle inequality. Consequently, several alternative metrics that more rigorously quantify similarity or dissimilarity between MSAs have been proposed (Blackburne and Whelan, 2012).

Machine learning (ML) has increasingly become a transformative tool in phylogenetics, enhancing both tree reconstruction and sequence alignment processes (Mo et al. 2024), as well as searching for computing branch support values (Ecker et al. 2024; Wiegert et al. 2024). For example, Leuchtenberger et al. (2020) employed neural networks to identify MSAs composed of four sequences that evolved under conditions of long-branch attraction or long-branch repulsion. Haag et al. (2022) used machine learning to quantify the “difficulty” of an MSA, i.e., to estimate how challenging it would be for a heuristic tree search to recover top-scoring tree topologies. BetaAlign, a deep learning-based method, utilizes transformer models and leverages natural language processing techniques to perform multiple sequence alignments (Dotan et al. 2024). However, this approach is currently limited in the number of sequences it can process effectively.

We introduce deep-learning–based approaches for improved scoring of MSAs and for ranking alternative alignments. Our approach involves generating diverse alternative MSAs using standard alignment tools, followed by evaluation and scoring with trained deep-learning models. We developed two types of models. *Model 1* provides an accuracy score for a single input MSA. Although it outperforms the SoP score, it does not perform well in identifying the top-ranking MSA from a set of alternatives. We therefore developed *Model 2*, which takes as input a set of alternative MSAs and predicts their relative ranking. We show that *Model 2* consistently identify MSAs that are significantly more accurate than those selected by the SOP metrics, *Model 1*, as well as MSAs produced by widely used tools such as MAFFT and PRANK. These findings are validated on both empirical and simulated datasets.

## Methods

To compute the SoP score, one must specify the substitution matrix and the affine gap penalty. In this paper, the default SoP score is calculated using the BLOSUM62 substitution matrix, with a gap opening penalty of -10 and a gap extension penalty of -0.5. These parameters were selected because they are the default settings in the EMBOSS Needle program, a widely popular pairwise alignment tool that employs affine gap costs (Rice et al. 2000). We use the fast algorithm suggested by Ranwez (2016) to compute this score efficiently. Of note, additional SoP scores with an additional substitution matrix and with different gap costs are computed as features (see description of the features used below).

We use the *d_seq_* measure to quantify dissimilarity between a pair of MSAs, because it is a proper mathematical distance metric (Blackburne and Whelan 2012). Let *A* denote an alignment of *n* sequences, and let *S^i^_j_* represent the *j*’th character of the unaligned sequence *S^i^*. Then, for a given sequence *S^i^* and alignment position *j*, we identify the alignment column in which *S^i^_j_* appears and define 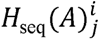 as the collection of all other characters (plus gaps if present) that occupy the same column. Thus, 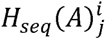 contains exactly *n*-1 elements and represents the set of characters (including gaps) that are homologous to *S^i^_j_* according to alignment *A*. Formally:

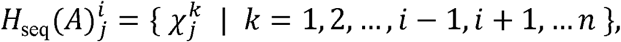

where

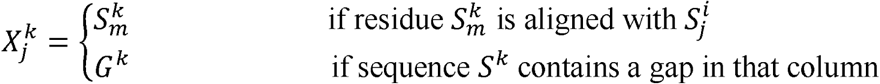

To illustrate, consider a toy alignment *A* of length two, with the following three sequences: ‘MC’, ‘MY’, and ‘M–’. For this example, 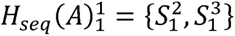, which represents the homology set of the first residue of the first sequence. Similarly, 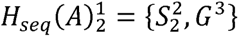. The contribution of the *j*’th position in sequence *S^i^* to the total distance between MSAs *A* and *B*, denoted *d_seq_*(*A,B)^i^_j_*, is defined as the dissimilarity between the set of characters aligned to *S^i^_j_* in the two alignments: 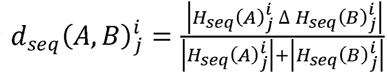, where Δ is the symmetric difference of two sets, i.e., the set of elements in either of two sets, but not in their intersection. The overall distance between the two alignments is computed as the average of these local dissimilarities across all characters: 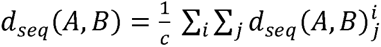, where *c* is the total number of characters across all sequences, i.e., the sum of all sequence lengths: *c* = ∑*_i_* |*S^i^*|.

We aimed to simulate MSAs that resemble empirical MSAs in terms of topology, branch lengths, model parameters (including indel and substitution rates), and alignment length. To this end, we first inferred these attributes from empirical data and used them to generate simulated MSAs. Empirical MSAs and their corresponding trees were extracted from OrthoMaM v.12, a database of orthologous mammalian protein-coding genes (Ranwez et al. 2007; Allio et al. 2024). We first selected 340 datasets, for which their corresponding phylogenetic trees exhibited the greatest total branch lengths. This criterion has been chosen because we expect that aligning highly variable sequences is more challenging. Subsequently, for each MSA, we randomly sampled a subset of either 40, 50, 60, 70, or 80 taxa (∼70 MSAs for each number of taxa). Each MSA is associated with the tree, and in each case, we pruned the tree to the corresponding taxa using the ETE3 package in Python (Huerta-Cepas et al. 2016). The tree branch lengths for OrthoMaM datasets were based on coding DNA alignments. Because we analyzed protein MSAs, we re-optimized the branch lengths of all the phylogenetic trees using IQ-TREE with *-s, -te,* and *-m LG+G8* parameters. The associated shape parameter of the gamma distribution (alpha) was extracted from the IQ-TREE log. We then inferred the indel parameters using SpartaABC v.0.6.0 (Ashkenazy et al. 2017). The program was executed with the following command: *sparta -i <PATH > -t AA -n 100000 -nc 300 -a mafft -s 12345*, and the *LG+G8{alpha}* model as indicated in the model.bestModel file.

Each of the resulting 340 datasets included a phylogenetic tree and the inference of substitution model parameters, indel model parameters, and the root sequence length. We then simulated 340 MSAs, referred to as “reference MSAs,” along with their corresponding unaligned sequence sets using INDELible v.1.03 (Fletcher and Yang 2009). These simulated data were subsequently used as reference datasets, which we denote as D_sim_.

We also benchmarked our algorithm against a set of empirical MSAs obtained from BAliBASE version 4 (Thompson et al. 2005). The benchmark included reference sets RV10, RV11, RV12, RV20, RV30, and RV50. These reference sets represent a wide variety of alignment challenges: reference set RV10 includes large, complex protein families and is specifically designed to address the challenges of the MSA construction algorithms while working with less frequent patterns; reference sets RV11 and RV12 contain divergent sequences with less than 20% and 40% identity respectively; reference set RV20 includes families aligned with a highly divergent “orphan” sequence; reference set RV30 contains subgroups with less than 25% residue identity between groups; reference set RV50 includes families where one or more sequences contain long inserted segments relative to the others. Hence, a total of 294 reference MSAs (“reference” empirical MSAs) were extracted from this dataset, which we denote as D_emp_.

To train and evaluate the deep-learning models, a large and diverse labeled dataset is required. Therefore, we generated 1,600 alternative alignments for each of the original “reference MSAs” derived from either the simulated or empirical datasets. The deep-learning algorithms will learn to score the accuracy of a given MSA (*Model 1*) or to rank the accuracy of a set of alternative alignments (*Model 2*). An MSA accuracy is defined as its distance from the corresponding reference alignment (see below). Each reference MSA with its associated inferred alternative MSAs is called hereafter an MSA-batch. The final dataset comprised a total of 1,600x340 = 544,000 and 1,600x 294 = 470,400 alternative MSAs for the simulated and the BAliBASE datasets, respectively.

The following steps were performed for each “reference” MSA to generate 1,600 alternative MSAs by realigning their sequences: (1) 800 alternative alignments were generated using GUIDANCE (Sela et al. 2015) v.3.1 with default parameters, 400 using MAFFT, and 400 using PRANK. GUIDANCE produces alternative alignments by introducing uncertainty in the guide trees and in the process of indel formation. (2) An additional 400 alternative MSAs were generated by Muscle5 (Edgar 2022) version 5.2.linux64, which perturbs the guide tree and the hidden Markov model (HMM) used for MSA construction and creates a so-called diversified ensemble of the alignments. Muscle alternative alignments were produced using the *-diversified-replicates 400-threads 8* parameters in the command line. (3) Another 400 alternative MSAs were extracted from the Markov Chain Monte Carlo (MCMC) iterations of each BAli-Phy (Gupta et al. 2021) version 3.6.1 run, which records a new alternative MSA every ten iterations. The BAli-Phy program was executed with the parameter *--iter=4000*.

For each reference MSA, we also obtained four “default MSAs,” defined as the standard inferred alignments produced by four different alignment programs using their default settings. Specifically, we first unaligned the sequences. We next realigned them using each of the following alignment algorithms: MAFFT (Katoh 2002; Katoh and Standley 2013) version v7.525, PRANK (Löytynoja 2014; Löytynoja 2021) v.170427, Muscle5 (Edgar 2022) version 5.2.linux64, and BAli-Phy (Gupta et al. 2021) version 3.6.1. The default alignments of MAFFT, PRANK, and Muscle5 were produced with the default settings. The BAli-Phy *bp-analyze* command line was run to produce *P1.max.fasta* alignment, which was used as the default alignment for this algorithm.

A total of 153 input features were extracted from each simulated MSA. Two additional features related to 3D protein structures were additionally extracted for empirical MSAs. The list of all 155 features is detailed in Supplemental Information S1. Each of the 155 features can be assigned to one of the following categories: (1) unaligned sequence attributes, e.g., the length of the unaligned shortest sequence; (2) MSA attributes, e.g., the total number of MSA columns; (3) SoP-related features, e.g., the sum of pairs cost, using the PAM250 substitution matrix with a gap opening and gap extension penalty of -10 and -1; (4) Gap-related features, e.g., the total number of gaps of length one in the MSA; (5) Tree-related features, e.g., the average branch length, when branch lengths are optimized on a neighbor-joining tree extracted from the MSA; (6) Entropy-related features, e.g., the average entropy of an alignment column; (7) k-mer related features, e.g., the average frequency of k-mer of length five found within the MSA; (8) Structure-related features, e.g., the fraction of columns that are identical between the MSA and a FoldMason-based structural alignment of the same set of sequences.

One of the primary objectives of this study was to develop a novel deep-learning-based scoring function for MSAs that correlates strongly with the quality of the inferred alignment, as measured by its *d_seq_* distance to the reference alignment. To this end, we trained a deep learning regression model that receives as input the MSA features and predicts *d_seq_,* referred to as *Model 1*. Supplemental Figure S1 illustrates the architecture of our neural network model, and Supplemental Table S1 provides additional network attributes and the optimal hyperparameters for this model. Hyperparameters were optimized using the Optuna package (Akiba et al. 2019), maximizing the correlation between the predicted and true *d_seq_*values. The above D_sim_ and D_emp_ data were each split into train, validation, and test sets in a 64:16:20 ratio. To avoid data leakage, all alternative MSAs derived from the same original dataset and based on the same sets of sequences were assigned to the same partition (train, validation, or test).

We subsequently trained a second model (referred to as *Model 2*), which incorporates modifications to *Model 1* that focus specifically on selecting the most accurate MSA from a given collection of alternatives. To support this objective, the target label was changed to a rank-based normalized version of the original *d_seq_* distance. Normalization was performed independently within an MSA-batch, i.e., the set of alternative MSAs derived from the same set of sequences. For a given alternative MSA with index *i* within an MSA-batch, the scaled target value was computed as: 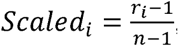, where *r_i_* represents the rank of the MSA with index *i* in this MSA-batch according to its *d_seq_* distance value, and *n* denotes the total number of alternative MSAs in the MSA-batch. In case of ties, all entries with the same rank were given the average ranks (e.g., if two MSAs had ranks 5 and 6 with the same score, both obtained the rank 5.5). Feature values were also normalized for all alternative MSAs within the same MSA-batch. For a given feature, all its feature values across the 1,600 alternative MSAs within an MSA-batch were ranked, and the scaled feature value was computed using the same rank-based approach described above for the target value.

When running *Model 2*, our objective is to distinguish between MSAs comprising an MSA-batch. During training and testing, a mini-batch of MSAs is processed by the deep-learning model (a mini-batch is a standard term in learning algorithms, which is the number of samples processed in each learning iteration. Note, this is a different “batch” than the MSA-batch). We implemented a custom “batch generator” that ensured all MSAs within a mini-batch belong to the same MSA-batch, i.e., *Model 2* “sees” each time it processes data only MSAs that belong to the same MSA-batch.

To emphasize the accurate ranking of the best alignments within each batch, the loss function was modified to penalize prediction errors more heavily for the top-performing MSAs. Specifically, the loss was defined as 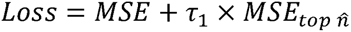, where MSE denotes the mean square error across the entire batch, and 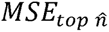 represents the mean squared error computed over the 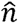 samples with the lowest (i.e., best) target labels within that batch. The free parameter τ_1_ controls the relative contribution of the MSE versus the 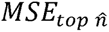 . The network attributes and hyperparameters of *Model 2* are listed in Supplemental Table S1. The selected hyperparameters maximized the validation loss. Both models were trained using TensorFlow Keras (Abadi et al. 2016; Chollet 2018).

All analyses were implemented in Python (v3.10). The source code, together with the OrthoMaM and BAliBASE dataset entries used in this study, is publicly available on GitHub (https://github.com/nimrodSerokTAU/sp_alternative). We provide detailed, step-by-step instructions for reproducing a representative example corresponding to MSA batch 126014, which was used to generate Figures 1b and 2b. This dataset contains 1,604 alternative MSAs, along with the corresponding true (simulated) MSA and the phylogenetic tree used to generate them.

**Figure 1.**
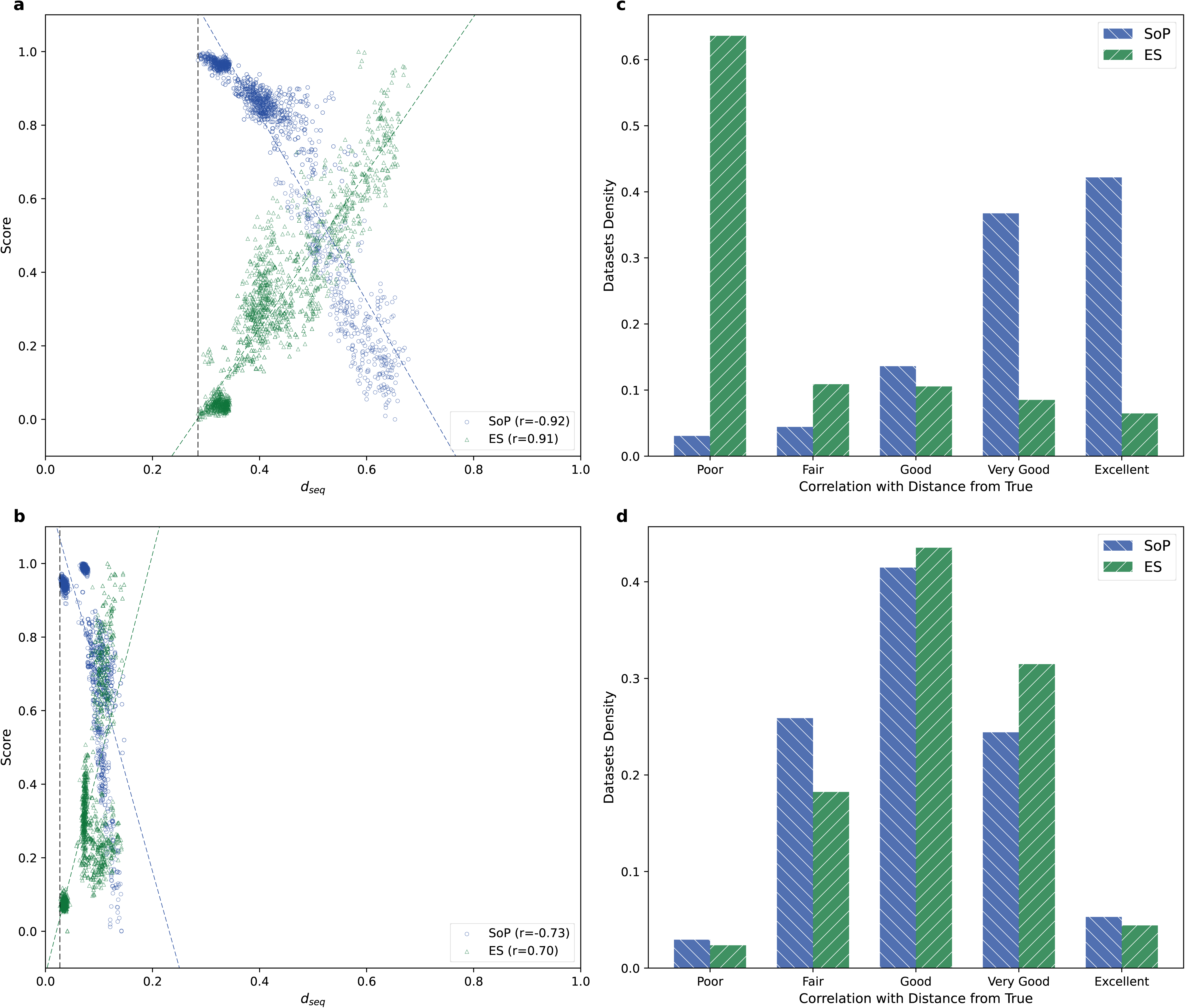
(a) Empirical MSA-batch BBA0058. Each dot represents the score obtained for one alternative MSA. For each alternative MSA we computed an alignment quality score (e.g., SoP) and *d_seq_* , the distance of the inferred MSA from the reference MSA. The Pearson correlation coefficient between each MSA quality score and *d_seq_* is shown at the lower right; (b) Similar analysis on a simulated MSA-batch; (c) Distribution of Pearson correlation coefficients across 294 empirical BAliBASE MSA-batches. Correlation strength for ES was categorized as follows: Excellent: (r ≥ 0.95), Very Good (0.85 ≤ r < 0.95), Good (0.70 ≤ r < 0.85), Fair (0.50 ≤ r < 0.70), and Poor (r < 0.50). For SoP, Excellent is defined as r ≤ -0.95 and similarly for the other categories; (d) Distribution of absolute values of Pearson correlation across 340 simulated MSA-batches, each reflecting the evolutionary dynamics of an OrthoMaM alignment.

**Figure 2.**
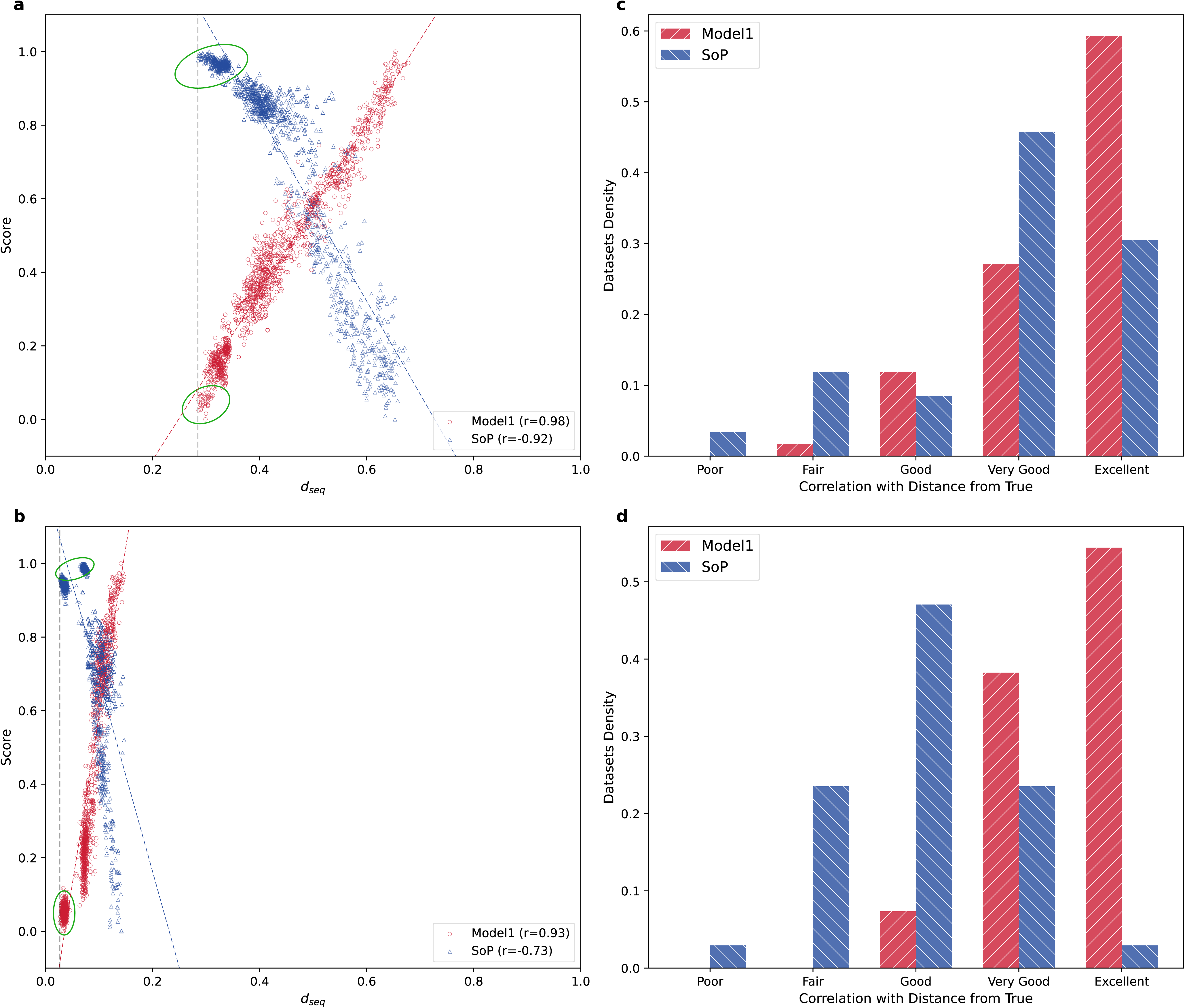
(a) Empirical MSA-batch BBA0058. The Pearson correlation coefficients between an MSA quality score (either SoP or *Model 1*) and *d_seq_* were computed. The quality scores were normalized to be between zero and one, using the formula 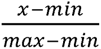, where *x*, *min* and *max*, correspond to the scores of the alternative MSA, the minimal score among all alternative MSAs, and the highest score, respectively. A vertical dashed line indicates the *d_seq_* value of the most accurate MSA among the alternatives. The oval shapes highlight top-scoring MSAs that both SoP and *Model 1* struggle to rank correctly; (b) Similar analysis on a simulated MSA-batch; (c) Distribution of absolute values of Pearson correlation across 59 empirical test MSA-batches for SoP and *Model 1*; (d) Similar analysis across 68 simulated test MSA-batches.

Our methodology takes as input a set of alternative MSAs. The first step consists of feature extraction across all MSAs in the batch. For the 1,604 MSAs in this example, feature extraction required 126 minutes and 35 seconds using a single CPU core. This step produces a CSV file summarizing the extracted features across all MSAs; this file is provided in the GitHub repository.

The feature file is then used as input to the trained deep neural network model. Model inference required less than one second and outputs a score for each MSA, provided as an additional CSV file. The source code used to generate all figures presented in this manuscript is also included in the repository.

Finally, this study is intended as a proof of concept demonstrating the feasibility of using machine learning to derive more accurate MSA scoring functions. Additional work will be required to translate this framework into an efficient, production-level tool. For example, feature extraction is currently implemented in Python and could be substantially accelerated through reimplementation in C++ and the use of parallel computing. Furthermore, most of the overall computational cost is devoted to generating the alternative MSAs used as input to our framework, a step that likewise presents substantial opportunities for optimization (see Discussion).

## Results

First, we aimed to assess the extent to which existing alignment quality measures are capable of predicting the distance from the reference alignment. Two alignment quality measures were considered: SoP and the entropy score (ES). The SoP quality measure is expected to be positively correlated with alignment quality (and negatively correlated with the distance from the reference MSA). The ES score is expected to be negatively correlated with alignment quality (and positively correlated with the distance from the reference MSA). To demonstrate how we evaluate the performance of each of these MSA quality metrics, consider a single MSA-batch, e.g., empirical MSA BBA0058 from the BAliBASE dataset. This MSA-batch includes a known reference alignment and 1,600 inferred alternative alignments, generated by various alignment tools. For each alternative MSA, we measured the alignment quality (e.g., the SoP score) and its distance from the reference alignment using the *d_seq_*distance metric, which has the advantage of being a mathematically valid distance measure that satisfies properties such as symmetry and the triangle inequality (Blackburne and Whelan 2012). We examined the Pearson correlation between the min-max normalized quality metric values and the distance from the reference alignment. For example, in the case of the SoP score, we expect higher values to correspond to smaller distances from the reference alignment, and vice versa. The Pearson correlation for this MSA-batch (BBA0058) was -0.92 (Figure 1a). The correlation coefficient for the ES was 0.91. Overall, SoP and ES performed comparably. We repeated the same analysis for a single simulated MSA-batch (Figure 1b). In this case, SoP performed slightly better than ES, with correlation coefficients of –0.73 and 0.70, respectively.

We next repeated this analysis across all 294 empirical BAliBASE MSA-batches (Figure 1c) and 340 simulated MSA-batches generated through a simulation process based on the OrthoMaM database (Figure 1d). On the empirical datasets, the SoP metric showed superior predictive accuracy compared to ES: the average correlation coefficients were -0.89 and 0.07 for SoP and ES, respectively. In contrast, for the simulated data, the ES performed slightly better: the average correlation coefficients were -0.76 and 0.79, for SoP and ES, respectively. Similar results were obtained when other MSA similarity metrics, namely Residues-Pairs and Toal Column, were used instead of the *d_seq_* metric (Supplemental Figure S2).

Using the above approach, we next evaluated our deep learning-based scoring scheme, hereafter referred to as *Model 1,* and compared its performance with that of the SoP metric on the same representative cases shown in Figure 1. For the empirical MSA-batch BBA0058, the Pearson correlation coefficients were 0.98 for *Model 1* and -0.92 for SoP (Figure 2a). *Model 1* also outperformed SoP on the simulated MSA-batch, with Pearson coefficients of 0.93 and -0.73, respectively (Figure 2b). Importantly, these test examples were not used during model training. A similar trend was observed when a larger set of test examples was analyzed (*Figure 2c, 2d*). As these panels display only the test data, excluding the training data, the number of datasets corresponds to one-fifth of the total: 59 empirical and 68 simulated datasets. Overall, these results indicate that across both empirical and simulated datasets, the two variants of *Model 1* (one for empirical and one for simulated data) consistently predict the distance from the reference alignment more accurately than the SoP metric.

*Model 1* predicts multiple MSAs close to the reference MSA, which appears as a scatter of dots in Figures 2a and 2b. These MSAs are characterized by similar small distances to the reference MSA. *Model 1* is unable to accurately identify the most accurate MSA out of this scatter. The same limitation is even more emphasized when the SoP metric is used.

This outcome is not surprising because the correlation calculation includes all evaluated alignments, including those that are very distant from the reference alignment, and *Model* 1 was trained implicitly assigning all of them with the same importance. Therefore, high overall performance across the entire dataset does not necessarily translate into accurate identification of the alignments closest to the reference MSA. However, researchers are often most interested in identifying the best alignment, rather than in accurately predicting distances for distant, low-quality alignments. In light of this, we proceeded to evaluate how well SoP and *Model 1* perform in the task of identifying the alignment closest to the reference alignment.

To further exemplify this issue, we analyzed the distribution of the distance from the reference MSA (*d_seq_*) over all 1,604 alternative and default MSAs, for a single empirical dataset (BBA0058) (Figure 3a). In this plot, the alternative MSA that was closest to the reference alignment has a *d_seq_* of 0.285. Note that this distance is non-zero, as the reference alignment itself was not among the alternative alignments generated by the inference tools. The *d_seq_* values associated with the default MSAs produced by the various aligners were 0.328, 0.329, 0.441, and 0.462 for MAFFT, Muscle, BAli-Phy, and PRANK, respectively. The inferred MSA with the highest SoP score yielded a *d_seq_* value of 0.306, while selecting the MSA with the lowest *Model 1* score resulted in a *d_seq_* value of 0.299. These findings highlight the limited effectiveness of *Model 1* in identifying the optimal MSA from a set of alternatives, despite its ability to predict with high accuracy the distance from the reference MSA when a large number of alternative MSAs are considered.

**Figure 3.**
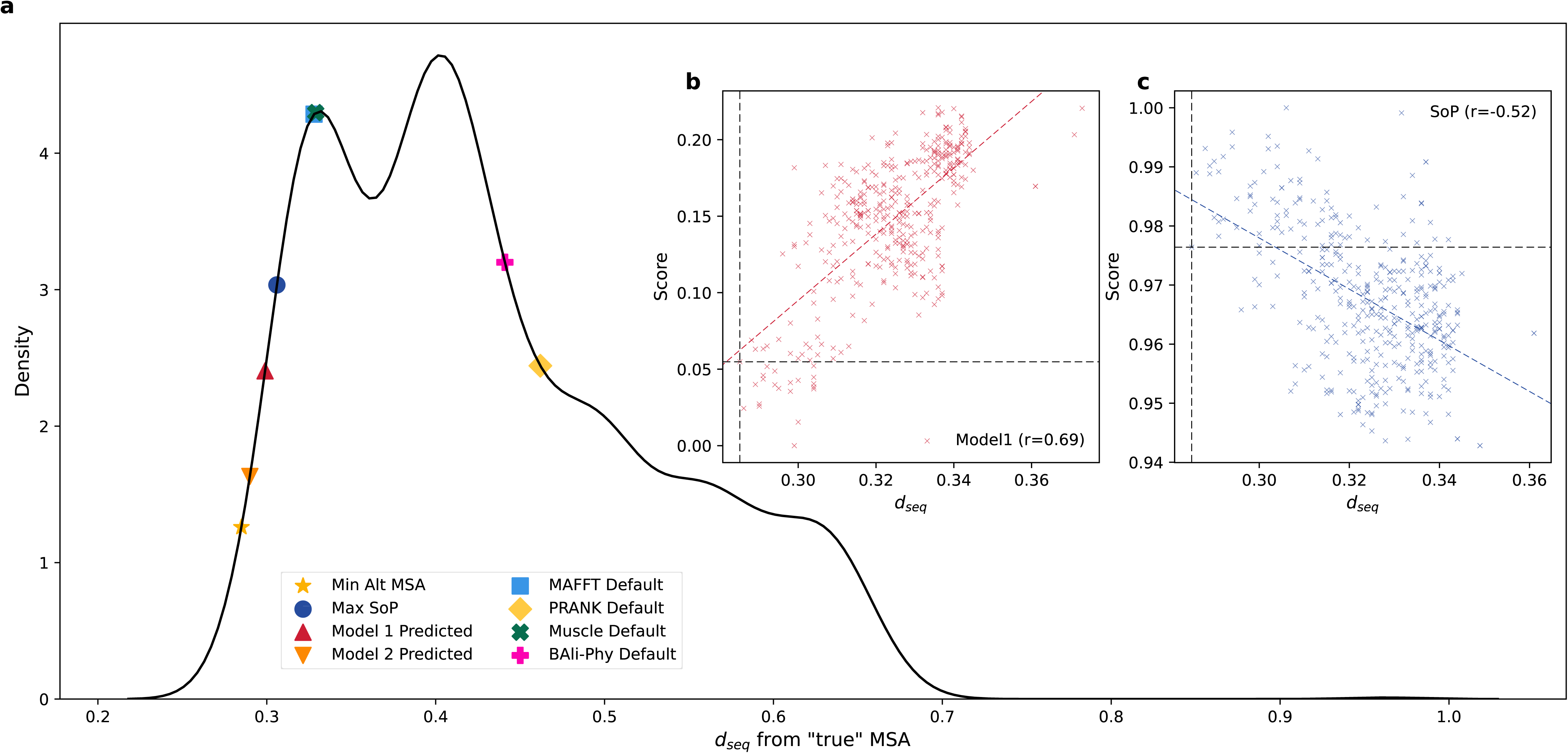
Analysis of empirical MSA-batch BBA0058. (a) Probability density function of *d_seq_* value for 1,604 alternative and default MSAs. Markers indicate the positions of the default alignments (MAFFT, PRANK, BAli-Phy, Muscle), the highest SoP-scoring MSA (“max SoP”), the MSA selected using Models 1 and 2, and the alternative MSA closest to the reference MSA (“Min Alt MSA”). The density observed to the left of the Min Alt MSA marker reflects a smoothing artifact introduced by the kernel density estimation (KDE) and does not correspond to additional alignments; (b) *d_seq_* and *Model 1* predicted scores for the 400 alternative MSAs of with the lowest *d_seq_*. The *d_seq_* and predicted score of the alternative MSA that is closest to the true MSA are indicated by vertical and horizontal lines, respectively. Alternative MSAs below the horizontal lines were predicted to be superior to the closest MSA, and hence represent clear errors of *Model 1*; (c) *d_seq_* and SoP predicted scores for the 400 alternative MSAs of with the lowest *d_seq_*. Alternative MSAs above the horizontal lines represent clear errors.

As stated above, among the alternative MSAs shown in Figure 3a, the one that was closest to the reference MSA had a *d_seq_* = 0.285. *Model 1* predicted the (normalized) score of this alternative MSA to be 0.0548. Among the 1,604 scores, 20 MSAs had predicted scores below this score (Figure 3b). When the goal is to find the optimal MSA, these are clear errors of *Model 1*. For comparison, using the SoP score, the (normalized) score of the closest MSA was 0.976, and 89 alternative MSAs had SoP values higher than this MSA (Figure 3c).

We next examined the relationship between the predicted score and *d_seq_* among the alternative MSAs closest to the reference MSA. Figure 3b focuses on the top 25% percent of alternative MSAs. As anticipated, the correlation observed within this subset (*r* = 0.69) is weaker than that observed across the full dataset (*r* = 0.98), highlighting *Model 1*’s limited ability to reliably rank MSAs that are close to the reference MSA. A similar pattern is observed for the SoP metric (Figure 3c), for which the correlation coefficient deteriorates from *r* = -0.92 for the entire data to *r* = -0.52 within the top 25%. These results underscore the need for an alternative score to discriminate among MSAs that are closest to the reference one.

To better discriminate between alternative MSAs that are similar to the reference one, we tested several alternatives, including: (1) Training on the subset of MSAs with the lowest *d_seq_*; (2) Training on augmented data of MSAs with the lowest *d_seq_*; (3) Alternative loss functions that penalize more heavily MSAs with low *d_seq_*; (4) Several ranking functions such as RankNet (Burges et al. 2005), Kendall loss (Kendall 1938; Brinker and Hüllermeier 2020), and NDCG (Chen et al. 2009). *Model* 2 (discussed below) outperformed these approaches, and hence we only show *Model* 2 results.

*Model 2* was trained to rank alternative MSAs and, accordingly, receives an MSA-batch as input. This is in contrast to *Model 1*, which gets as input a single MSA and predicts its accuracy. *Model 2* was trained using ranked-normalized target values, and its loss function penalizes errors in the relative ranking of accurate MSAs more heavily (see Methods). For the example analyzed in Figure 3, *Model 2* selected an inferred MSA with *d_seq_*= 0.29, better than the MSA predicted by *Model 1* (*d_seq_* = 0.299) and only 1.8% greater distance than the best possible alignment (*d_seq_* = 0.285). The prediction of *Model 2* was better than those selected by all default aligners. Specifically, the distance from the best alternative alignment was 13% to 59% higher for the default aligners compared to *Model 2* alignments (Figure 3a). We next tested the hypothesis that *Model 2* outperforms *Model 1* in terms of correlation with *d_seq_*. This was not the case (Supplemental Figure S3), highlighting the differences between the two models: while *Model 1* can predict MSA quality without requiring alternative MSAs, *Model 2* is trained to pick the top-scoring MSA among alternative MSAs, a task it performs better than *Model 1*.

We next evaluated the performance of *Model 2* on a large set of datasets using the “pick best MSA” task, designed to quantify the ability to identify the most accurate alignment from a set of alternative MSAs. In each instance, the model was tasked with selecting a single alignment, and this choice was compared against selections made by the SoP score and the default outputs of the alignment algorithms. As shown in Figure 4a, *Model 2* selected alignments that were closer to the reference alignment than those chosen by SoP in 69.5% and 82.4% of the cases in empirical and simulated sets, respectively (P < 0.001, Chi-square test with df = 1, in both cases). Except for the case of BAli-Phy, *Model 2* also outperformed the default aligners, consistently choosing better MSAs, both for empirical and simulated datasets (Figure 4 b, c, respectively). These results underscore the practical effectiveness and value of *Model 2* in real-world scenarios and demonstrate its superiority over both traditional scoring methods and alignment tool outputs.

**Figure 4.**
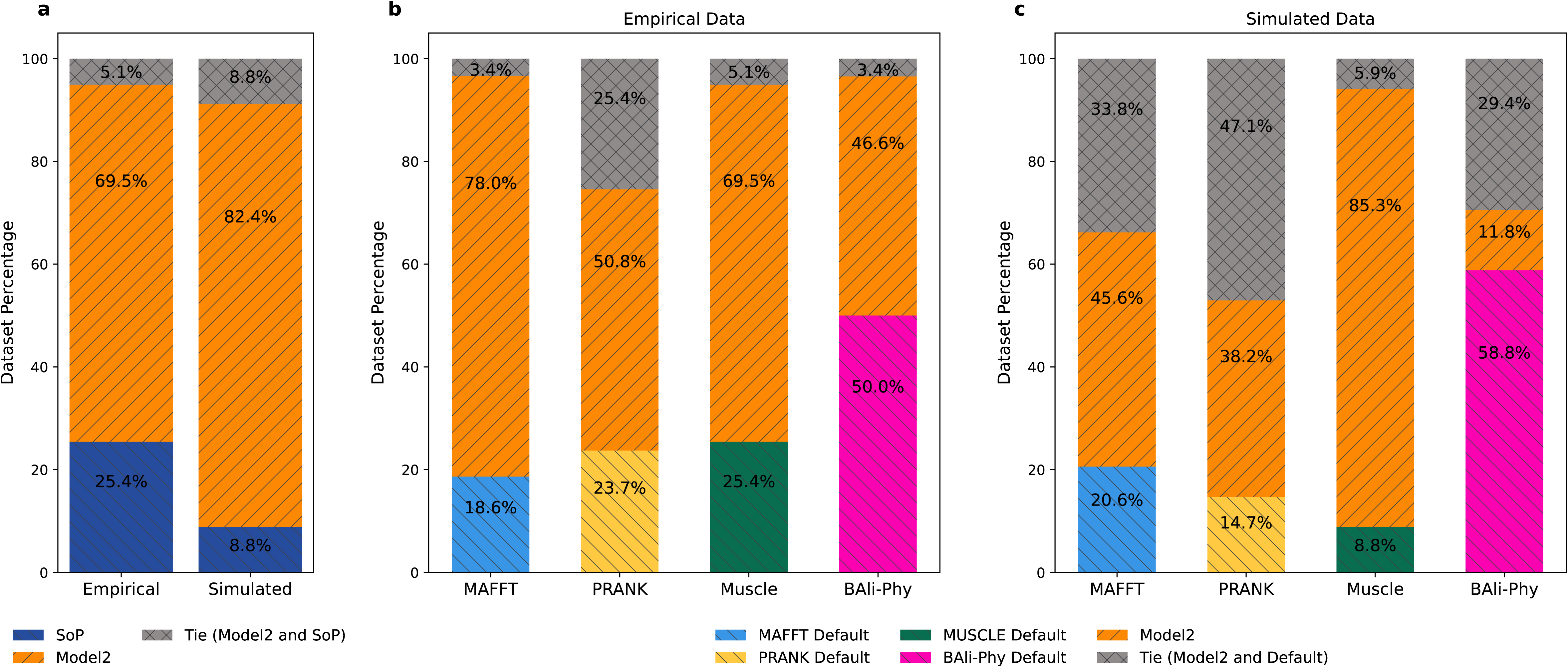
Performance of *Model 2* on the “pick best MSA” task. For each MSA batch, each method (*Model 2*, the SoP score, or an aligner’s default output) selected a single MSA from the set of alternative alignments, and performance was assessed by comparing the distance of the selected MSA from the reference MSA. (a) Proportion of MSA batches in which *Model 2* or the SoP score identified the more accurate alignment, or in which both methods tied; shown separately for the empirical (59 batches) and simulated (68 batches) datasets. (b) For the empirical dataset, proportion of cases, stratified by alignment program, in which *Model 2* selected a more accurate MSA than the default output of that program, the default output was more accurate, or the selections tied. (c) Same as panel (b), but for the simulated test set.

To gain insights into the decision-making process of our models, we analyzed the 40 most important features used by regression-based *Model 1* and ranking-optimized *Model 2*. We assessed features importance using SHAP values (SHapley Additive exPlanations) (Lundberg and Lee 2017), which quantify the contribution of each feature to the final prediction (Table 1). We found noticeable differences between features important for *Model 1* and for *Model 2*, reflecting their distinct training objectives. In *Model 2*, we observed a strong enrichment of SoP-based features, which include SoP scores calculated using different substitution matrices (e.g., BLOSUM50, BLOSUM62, PAM250) and various gap opening and gap extension penalty combinations. This diversity suggests that no single parameter set is universally optimal across all datasets, and that the model can benefit from using multiple sets of parameters. Rather than relying on a fixed configuration, *Model 2* leverages a broader landscape of scoring regimes simultaneously, which allows it to adapt to data-specific characteristics. This observation highlights one of the major limitations of the non-BAli-Phy approaches that rely on a single fixed set of parameters. In addition to SoP variants, both models consistently assign high importance to structural and statistical features such as entropy, total branch lengths, number of unique gap patterns, parsimony statistics, and k-mer-based similarity metrics. These features clearly capture complementary characteristics of MSA quality that are not captured by SoP alone.

**Table 1.**
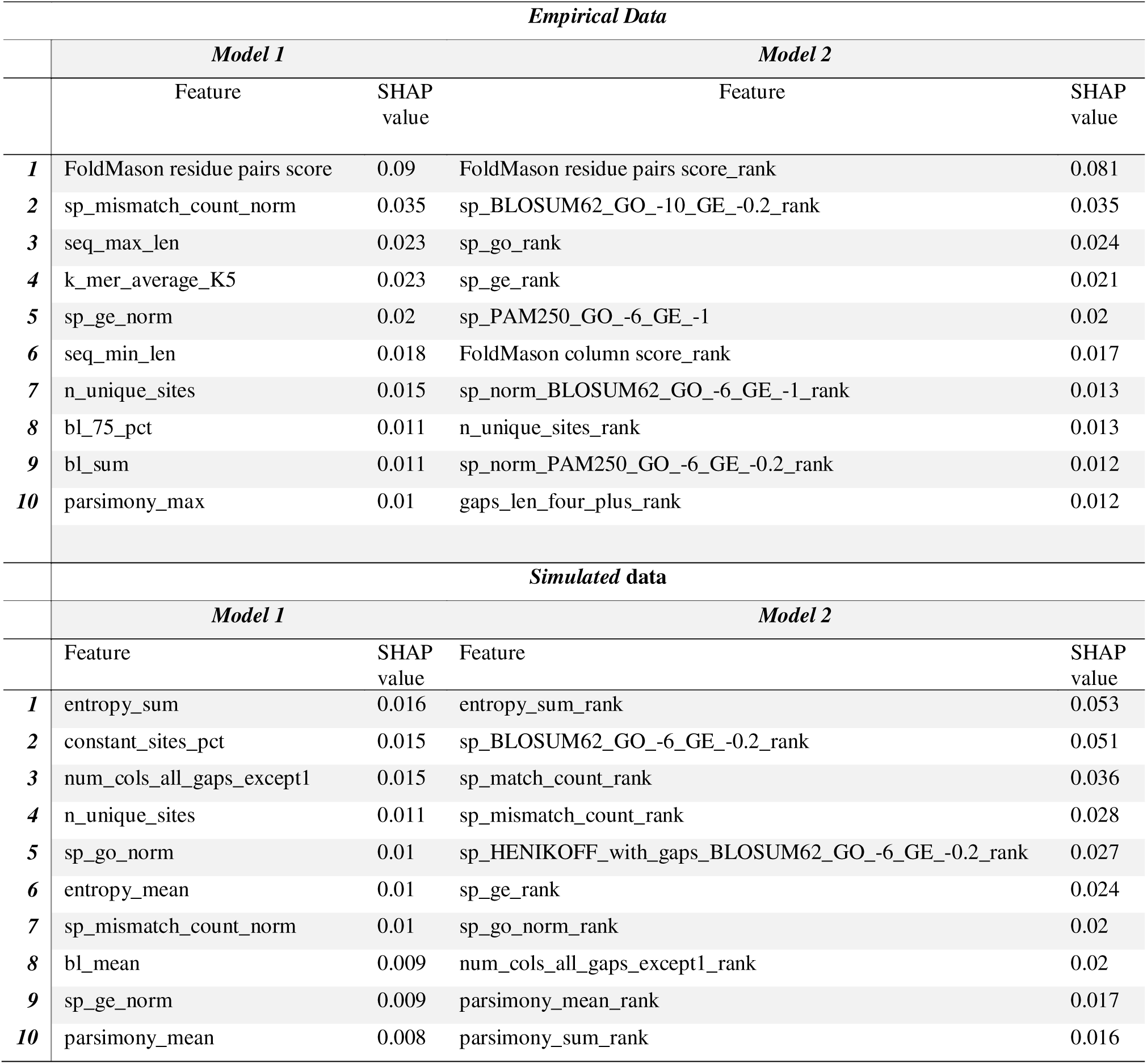
features importance using SHAP values, which quantify the contribution of each feature to the final prediction.

To evaluate the effect of the underlying MSA on phylogenetic accuracy, we compared the normalized Robinson–Foulds (RF) distances between inferred and true trees across 68 test simulated datasets. Trees reconstructed from the reference MSAs and those from the MSAs with the best *Model 2* predicted scores exhibited very similar mean RF distances (0.293 vs. 0.300, respectively; *p-value* = 0.124, two-sample t-test). In contrast, trees reconstructed from MAFFT default MSAs exhibited significantly reduced topological accuracy compared to those inferred from the best predicted MSAs (mean RF distance of 0.321 vs. 0.293; *p-value* = 5.1×10□□). Trees inferred from the MSAs generated using the default MUSCLE and PRANK aligners exhibited even lower RF distance than those inferred from MAFFT (not shown). Overall, these results demonstrate that phylogenies inferred from MSAs selected using the trained *Model 2* ranking function achieved accuracy comparable to those based on the true alignments, and outperformed trees reconstructed from standard MAFFT default alignments.

## Discussion

In this work, we developed two deep learning models that address distinct aspects of MSA evaluation. *Model 1* acts as a scoring function, mapping any given MSA to a single numerical value that serves as a proxy for its likelihood or overall quality. *Model 2*, in contrast, directly compares alternative MSAs. In principle, a scoring function can also be used to rank alignments by sorting their predicted scores. However, our results show that while *Model 1* correlates strongly with overall alignment quality, it did not perform well in distinguishing among top-scoring MSAs. This limitation likely arises because *Model 1* was not explicitly trained to discriminate between closely related, high-quality alignments. Consequently, the correlation between predicted and true quality decreases substantially when only top-ranked MSAs are considered. These observations motivated the development of *Model 2*, which uses a dedicated loss function that optimizes relative ordering among MSAs and assigns a higher weight to MSAs that are close to the reference alignment. This approach reflects the practical goal of most applications: to identify a single high-quality MSA suitable for downstream analyses, rather than to assign absolute scores to all possible alignments. A key innovation of our framework, therefore, is that unlike SoP and related metrics, *Model 2* explicitly optimizes for ranking accuracy rather than for absolute scoring.

To our knowledge, this study is the first to employ deep learning to learn an MSA ranking function. A major distinction between our approach and classic scoring schemes such as SoP and its variants is that we use supervised learning to directly train the ranking model. This allows the model to capture complex dependencies among features, thus integrating over multiple characteristics of an MSA and not just the inventory of the number of gaps and substitutions. Machine learning methods are increasingly being applied to algorithmic challenges in molecular phylogenetics (Azouri et al. 2021; Smith and Hahn 2023; Azouri et al. 2024; Mo et al. 2024). Here, we demonstrate that such approaches can also address one of the field’s most persistent problems, MSA inference. Remarkably, our trained model already produces alignments that are, on average, more accurate than those generated by widely used programs such as MAFFT, PRANK, and MUSCLE 5, both on simulated and empirical datasets.

We designed over 150 handcrafted features that capture diverse structural and statistical properties of both the MSA and its underlying phylogenetic tree. These features allow our models to overcome several limitations of traditional scoring methods, such as SoP, which depend on a single affine-gap and substitution model to evaluate pairwise alignments. Many of our features generalize this concept by varying substitution matrices and gap penalties. For example, Feature 27 (“sp_BLOSUM62_GO_-6_GE_-0.2”) represents the SoP score computed using the BLOSUM62 matrix with a gap opening penalty of −6 and a gap extension penalty of −0.2, whereas Feature 37 (“sp_PAM250_GO_-10_GE_-1”) uses the PAM250 matrix with a gap opening penalty of −10 and extension penalty of −1 (see Supplemental Information S1 for details). By learning dependencies among such features, the model can adaptively downweigh Feature 27 when few long indels are present, while assigning greater importance to Feature 37. The model learns whether there are few or many long indels, from features such as “gaps_len_four_plus,” which counts gaps of length ≥4, and “avg_unique_gap_length,” which quantifies the average length of unique indels. As part of feature extraction, we also compute a neighbor-joining tree and calculate parsimony scores for each alignment column, e.g., “parsimony_mean,” which measures the mean parsimony score across all columns. Together, our features capture both parsimony-based concepts underlying early alignment algorithms and SoP-based measures that characterize most modern MSA inference methods.

In this study, we trained each model twice: once using simulated datasets and once using empirical datasets. We observed that models trained on simulated data performed less effectively when applied to empirical data, and vice versa. Additionally, *Model 2* outperformed all aligners, except BAli-Phy. These findings are consistent with previous studies demonstrating fundamental differences between simulated and empirical datasets. For example, Trost et al. (2024) showed that a machine learning classifier can readily distinguish between empirical and simulated MSAs. Another important distinction between the simulation-trained and empirically trained models lies in the features used during training. The empirical models incorporated structure-based features, which are irrelevant to the simulation-trained models because the simulated datasets did not account for protein structural constraints. Finally, some of the apparent prediction errors observed for the empirical models may stem from inaccuracies in the empirical reference alignments themselves. Unlike simulated MSAs, which have a known ground truth, the reference alignments in datasets such as BAliBASE are inferred and therefore cannot be regarded as genuine reference MSAs (Edgar 2010; Iantorno et al. 2014). Despite these differences between models trained on simulated versus empirical MSAs, our results clearly show that accounting for multiple features capturing global aspects of the MSA within a deep learning inference framework can be highly efficient in ranking both types of alternative MSAs and thus help improve the accuracy of MSA inference.

We have shown that *Model 2* enables selecting a single MSA that is more accurate than several widely used alignment programs. From the practical aspect, however, this necessitates generating alternative MSAs and running the trained model on each of them to select the best one, which is computationally costly. Relatively straightforward modifications can potentially reduce running times, e.g., discarding some of the alternative MSAs based on a subset of features and extracting the entire set of informative features only from those alternative MSAs that pass the first filtering step.

Additional types of features could further enhance model performance. For instance, when analyzing empirical data, one could incorporate features derived from predicted three-dimensional protein structures generated by tools such as AlphaFold3 (Abramson et al. 2024), though this approach would incur substantial computational cost. A promising alternative is to use embedding vectors from protein language models, which capture rich contextual information about amino acid sequences (Rao et al. 2021; Rives et al. 2021; Brandes et al. 2022; Lin et al. 2023; Heinzinger et al. 2024; Hayes et al. 2025). Because several features in our current approach depend on the assumed phylogenetic tree, a fully end-to-end framework would also require direct embedding of the tree. Nevertheless, handcrafted features retain important advantages in interpretability. They facilitate the identification of systematic failure modes and provide biological insight into model behavior. For example, in cases of long-branch attraction (Susko and Roger 2021), deviations in tree-based features can reveal the source of alignment error, an effect that might be obscured when using purely learned representations.

For training the deep learning models, we applied fully connected networks with a handful of hidden layers. The loss function for *Model 2* was relatively ad-hoc, aiming to give more weight to erroneous ranking of alternative MSAs that are close to the reference one than the rest of the alternative alignments. A few networks dedicated to ranking were previously suggested (Burges et al. 2005; Chen et al. 2009; Brinker and Hüllermeier 2020b; He et al. 2022). In addition, transformers were shown to be a promising architecture for models that use representation learning. We predict that a substantial increase in model performance can be obtained by exploring alternative network configurations and loss functions, following innovations in the very fast-developing field of AI research.

Further research is required to fully characterize and optimize the proposed methodology. For instance, when training an MSA ranking model, it remains unclear whether performance would benefit more from increasing the number of alternative MSAs within each MSA-batch or from expanding the total number of MSA-batches. Another open question concerns dataset composition: should “easy” cases, such as alignments with low indel rates, be excluded from the training and validation sets to focus the model on more challenging examples? Furthermore, we generated alternative MSAs in equal proportions using the alignment tools PRANK, MAFFT, MUSCLE, and BAli-Phy. It is possible, however, that some alignment methods should be more heavily represented within each training batch than others, depending on their error characteristics or similarity to the true alignment. Systematic exploration of these design parameters will be valuable for improving both training efficiency and model generalizability.

*Model 2* currently ranks a limited set of alternative MSAs for each group of unaligned sequences, leaving many potentially high-scoring alignments unexplored. In this study, alternative MSAs were generated, for example, using GUIDANCE2 (Sela et al. 2015) by perturbing guide trees, adjusting gap penalties, and sampling co-optimal solutions within progressive alignment frameworks. Each resulting MSA therefore represents a distinct combination of these alignment parameters. This procedure could be extended into an iterative refinement framework. Specifically, one could begin by sampling alternative MSAs, use *Model 2* to identify the top candidates, and then generate new alternatives by applying targeted perturbations to these top-performing MSAs (e.g., small modifications to guide trees or gap penalties). This cycle would iteratively refine the candidate pool while balancing exploration and exploitation. Additional promising MSAs might also be generated by combining information from several top-ranking alignments (Collingridge and Kelly 2012). Alternative refinement algorithms can also be used to search for neighboring MSAs with a high *Model 2* score (Gotoh 1996; Thompson et al. 2003; Gotoh 2014; Katoh and Standley 2014; Mokaddem et al. 2019). Such iterative processes terminate once no additional high-scoring MSAs are found.

One limitation of our simulation-based *Model 2* is that it is trained and tested on MSAs generated under indel and substitution stochastic models that fail to capture several key aspects of molecular sequence evolution. For instance, the substitution process assumes a constant rate within each site, even though it is well established that evolutionary rates can vary over time at the same site, a phenomenon known as heterotachy (Lopez et al. 2002; Galtier and Jean-Marie 2004). Similarly, given the tree topology and site-specific rates, our simulator models substitutions at different alignment positions as independent events, thereby ignoring potential co-evolution between residues. When the three-dimensional structure of a protein is known, incorporating structural constraints into the simulations would provide a more biologically realistic representation of sequence evolution (Robinson et al. 2003; Rodrigue et al. 2005; Echave and Wilke 2017; Ferreiro et al. 2026). In addition, although the indel model used here allows for distinct insertion and deletion rates (Loewenthal et al. 2021), it overlooks several well-documented aspects of indel evolutionary dynamics (Redelings et al. 2024). For example, the current model assumes that indel probability is uniform across all alignment positions, implying no among-site variation in indel rate. This assumption is clearly oversimplified, as indels are typically rare in the structurally constrained core of proteins and more frequent on protein surfaces (Kim and Guo 2010). Overall, increasing the biological realism of simulated MSAs represents an important direction for future work. Such improvements are likely to enhance not only the accuracy of deep-learning-based MSA inference but also the performance of downstream deep-learning-based analyses in molecular evolution, including phylogenetic tree reconstruction, estimation of phylodynamic model parameters (Voznica et al. 2022; Lambert et al. 2023), and model selection (Abadi et al. 2020; Burgstaller-Muehlbacher et al. 2023; Kulikov et al. 2024; Braichenko et al. 2025).

In this work, we focused on identifying a single MSA that best represents the underlying homology relationships among the analyzed sequences. However, when the ultimate goal is phylogenetic reconstruction, it is often advantageous to account for alignment uncertainty rather than relying on a single, fixed MSA (Wheeler et al. 1995; Sela et al. 2015). More generally, the Bayesian framework for phylogenetic inference advocates integrating, or averaging, over sources of uncertainty, such as model parameters and alignment uncertainty, rather than conditioning on point estimates (Lunter et al. 2005; Hohna et al. 2016; Baele et al. 2017; Nascimento et al. 2017). A promising future direction would be to develop deep-learning models capable of generating samples of alternative alignments drawn according to their posterior probabilities. For downstream inference tasks that depend on alignments as intermediate representations, an even more ambitious goal would be to bypass explicit alignment altogether by enabling deep-learning models to process unaligned sequences directly.

## Acknowledgments

K.P. and N.S. were supported by fellowships from the Edmond J. Safra Center for Bioinformatics at Tel Aviv University. K.P was also supported by the Khazanov scholarship program and the TAD Excellence Program for Doctoral Students in Artificial Intelligence and Data Science. T.P. was supported by Israel Science Foundation grant 2818/21.

