## Supplementary material for "A deep-learning-based score to evaluate multiple sequence alignments": SOP_sup_mateial_1.docx

Nimrod Serok^1,*^, Ksenia Polonsky^1,*^, Haim Ashkenazy^2^, Itay Mayrose^3^, Jeffrey L. Thorne^4,5^, Tal Pupko^1†^

^1^ [The Shmunis School of Biomedicine and Cancer Research](https://en-lifesci.tau.ac.il/lp-en-mcbb), George S. Wise Faculty of Life Sciences, Tel Aviv University, Tel Aviv 69978, Israel.

^2^ Department of Molecular Biology, Max Planck Institute for Biology Tübingen, Tübingen, Germany.

^3^ The School of Plant Sciences and Food Security, George S. Wise Faculty of Life Sciences, Tel Aviv University, Tel Aviv 69978, Israel

^4^ Department of Biological Sciences, North Carolina State University, Raleigh, NC 27695, USA.

^5^ Department of Statistics, North Carolina State University, Raleigh, NC 27695, USA.

*These two authors equally contributed to this work

**Supplemental Table S1**. Hyperparameter and architectural specifications for the four trained models. The table summarizes all model configurations used in this study.

|  | Simulated | | Empirical | |  |
| --- | --- | --- | --- | --- | --- |
|  | *Model 1* | *Model 2* | *Model 1* | *Model 2* | Ref and remarks |
| Number of layers | 4 | 4 | 4 | 3 |  |
| Number of neurons in each layer | 256, 16, 128, 64 | 64, 128, 64, 512 | 64, 64, 512, 128 | 190, 180, 256 |  |
| Activation function | PReLU | | | | (He et al. 2015) |
| Output layer | Linear | | | |  |
| Normalization | Batch normalization | | | | (Ioffe and Szegedy 2015) |
| Dropout rate | 0.34 | 0.24 | 0.10 | 0.32 |  |
| Regularization | L2, strength parameter set to $6.77\times{10}^{-5}$ | L2,  strength parameters set to 1$.65\times{10}^{-5}$ | L1-L2 Elastic Net  strength parameters set to $2.77\times{10}^{-6}$ and $1.03\times{10}^{-5}$ | L1-L2 Elastic Net  strength parameters set to $2.83\times{10}^{-5}$ and 4$.16\times{10}^{-7}$ | (Zou and Hastie 2005) |
| Optimizer | ADAM | | | | (Kingma and Ba 2015) |
| Initial learning rate | 0.0001 | 0.0022 | $0.00009$ | 0.0022 |  |
| Callbacks | Learning rate scheduler, early stopping | | | |  |
| Mini-batch size | 128 | 32 | 64 | 64 |  |
| Maximum number of training epochs | 50 | | | |  |
| $\tau_{1}, \hat{n}$ | N/A | 1.33, 8 | N/A | 0.28, 8 |  |

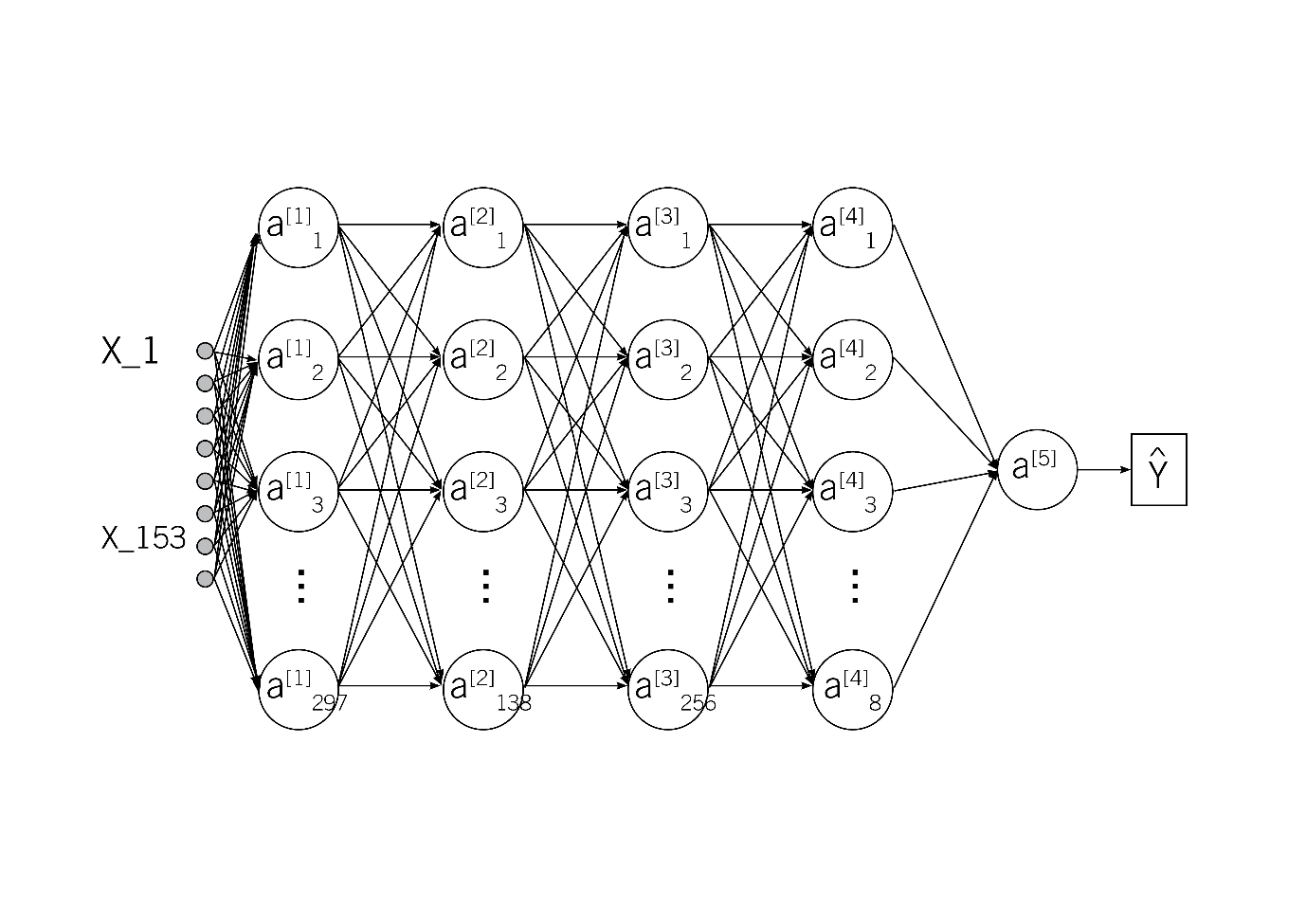
Figure S1. A fully connected deep network architecture for *Model 1*. The number of nodes in the hidden layer equals the number of features (153 when analyzing simulated MSAs). The last layer is a sigmoid activation function that provides values between zero and one, predicting the distance between two MSAs according to $d_{seq}$. The number of layers, nodes in each layer and additional hyperparameters were optimized based on the validation dataset.

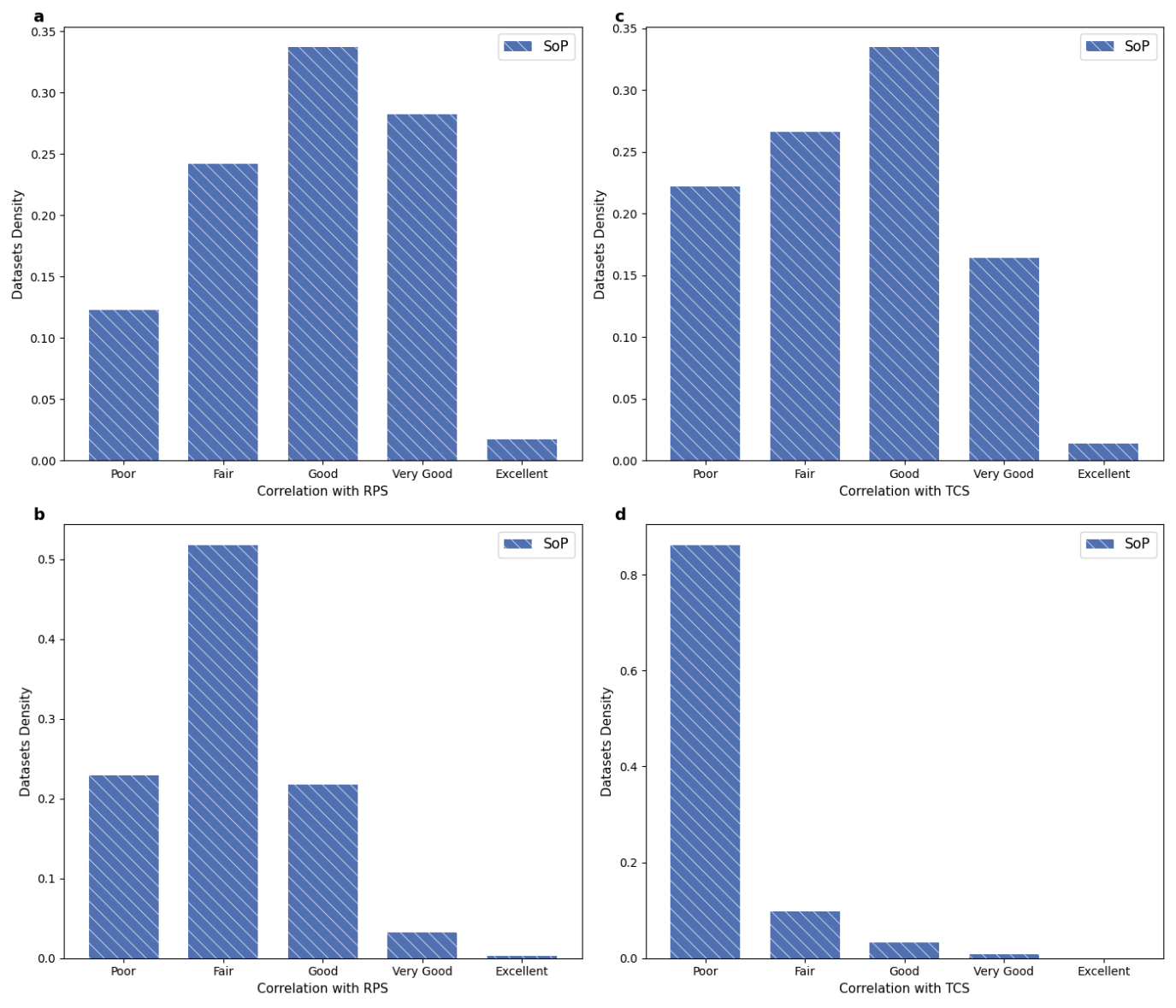

Figure S2. (a) Distribution of Pearson correlation coefficients of SoP metric and RPS metric (from reference MSA) across 294 empirical BAliBASE MSA-batches. Correlation strength was categorized as follows: Excellent (r ≥ 0.95), Very Good (0.85 ≤ r < 0.95), Good (0.70 ≤ r < 0.85), Fair (0.50 ≤ r < 0.70), and Poor (r < 0.50); (b) Distribution of Pearson correlation coefficients of SoP metric and RPS metric across 340 simulated MSA-batches, each reflecting the evolutionary dynamics of an OrthoMaM alignment. (c) Similar to (a), but the metric is TCS score (from reference MSA). (d) Similar to (b), but the metric is TCS score.

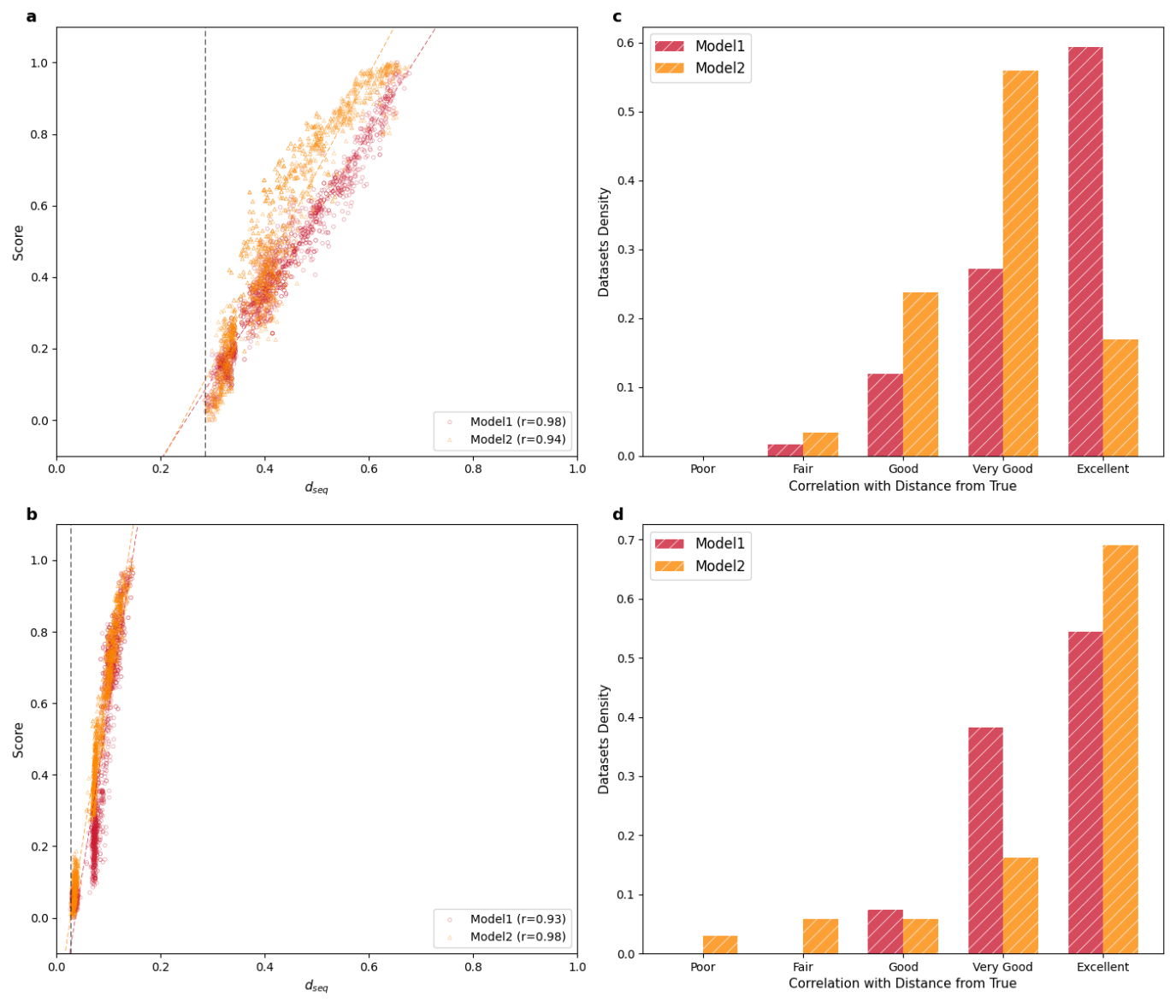

Figure S3. (a) Empirical MSA-batch BBA0058. The Pearson correlation coefficients between an MSA quality score predictors *Model 1* and *Model 2*, and $d_{seq}$ were computed. The quality scores were normalized to be between zero and one, using the formula $\frac{x-min}{max-min}$, where $x$, $min$, and $max$, correspond to the scores of the alternative MSA, the minimal score among all alternative MSAs, and the highest score, respectively. A vertical dashed line indicates the $d_{seq}$ of the most accurate MSA among the alternatives; (b) Similar analysis on a simulated MSA-batch; (c) Distribution of Pearson correlation coefficients across 59 empirical MSA-batches for *Model 1* and *Model 2* (average $r$ values 0.93 and 0.88, respectively); (d) Similar analysis across 68 simulated MSA-batches (average $r$ values 0.94 and 0.92 for *Model 1* and *Model 2*, respectively).
