## Supplementary figures and images for "A deep-learning-based score to evaluate multiple sequence alignments"

### FigS2_294_294_340_340.pdf

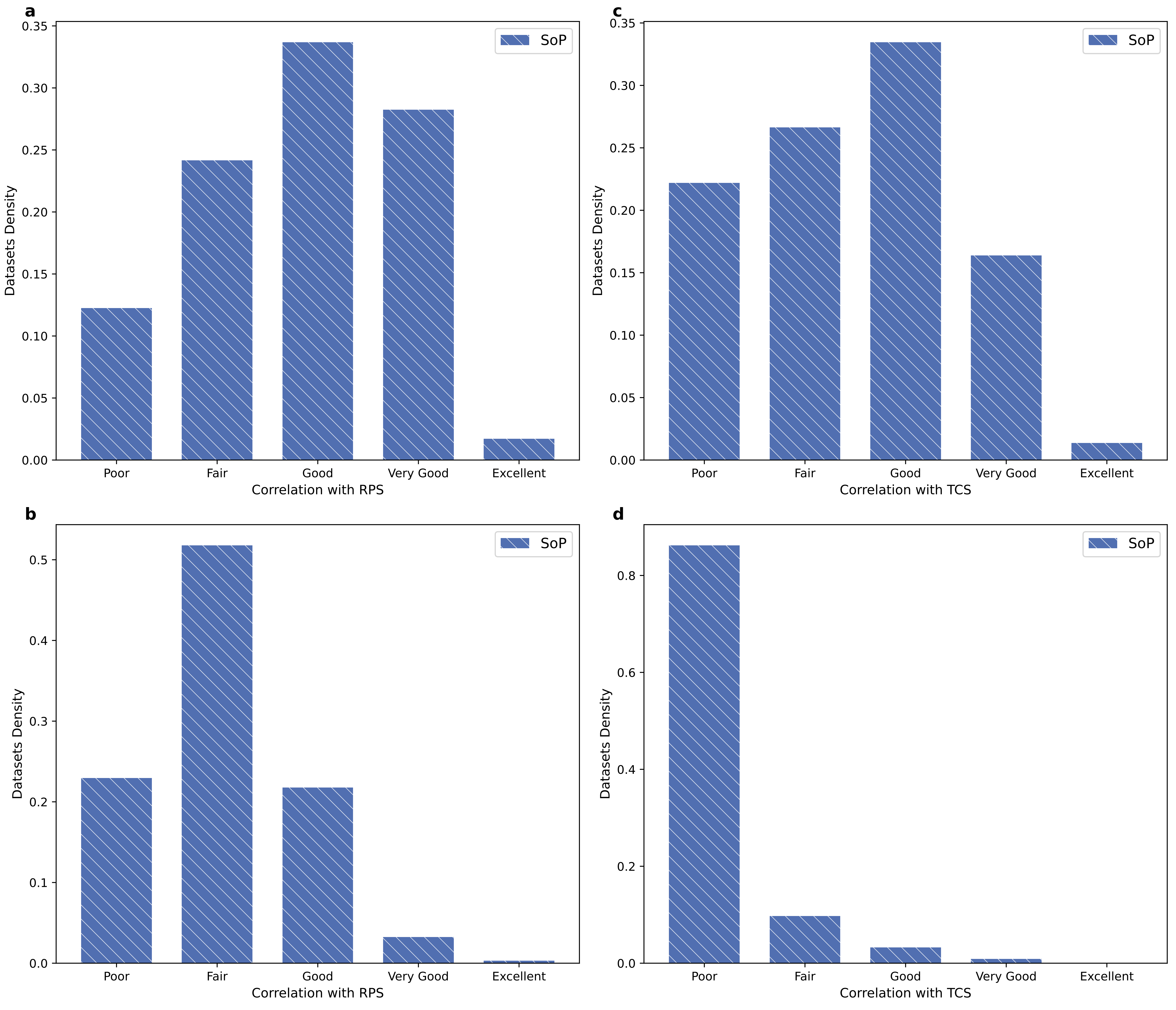

### FigS3_1603_59_1603_68.pdf

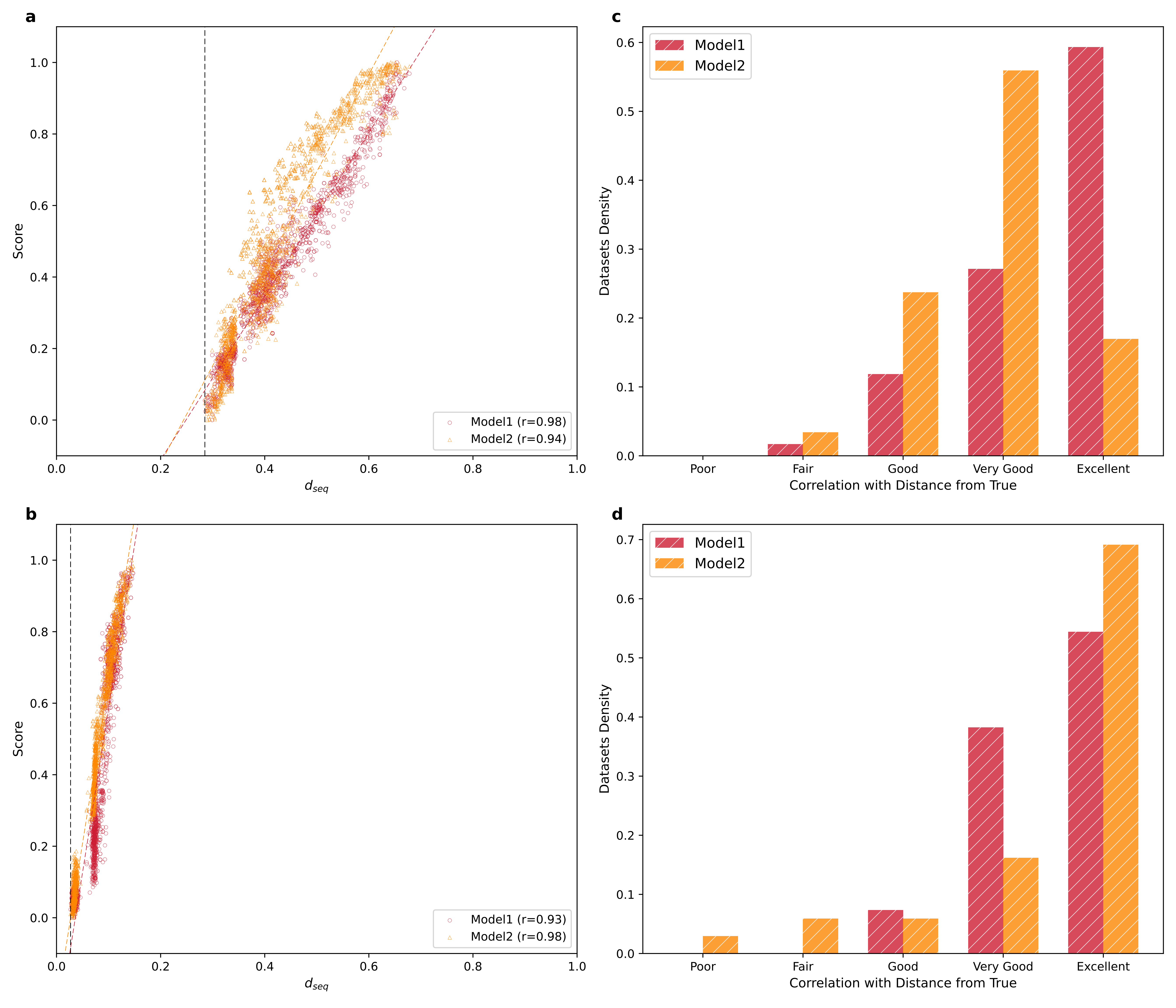
