## Supplementary material for "A deep-learning-based score to evaluate multiple sequence alignments": SOP_sup_information.docx

**Supplementary Information**

Nimrod Serok^1,*^, Ksenia Polonsky^1,*^, Haim Ashkenazy^2^, Itay Mayrose^3^, Jeffrey L. Thorne^4,5^, Tal Pupko^1†^

^1^ [The Shmunis School of Biomedicine and Cancer Research](https://en-lifesci.tau.ac.il/lp-en-mcbb), George S. Wise Faculty of Life Sciences, Tel Aviv University, Tel Aviv 69978, Israel.

^2^ Department of Molecular Biology, Max Planck Institute for Biology Tübingen, Tübingen, Germany.

^3^ The School of Plant Sciences and Food Security, George S. Wise Faculty of Life Sciences, Tel Aviv University, Tel Aviv 69978, Israel

^4^ Department of Biological Sciences, North Carolina State University, Raleigh, NC 27695, USA.

^5^ Department of Statistics, North Carolina State University, Raleigh, NC 27695, USA.

*These two authors equally contributed to this work

### Supplemental Information S1

**Features:**

**1. Unaligned sequence attributes**

Feature 1. Name: “seq_max_len”. The length on the unaligned (not including gaps) longest sequence.

Feature 2. Name: “seq_min_len”. The length on the unaligned shortest sequence.

**2. MSA attributes**

Feature 3. Name: “msa_lengh”. Let *L* denote the MSA length. This is simply the number of MSA columns.

Feature 4. Name: “taxa_num”. Let *N* denote the number of sequences in the MSA. This is the number of sequences in the MSA.

**3. SoP-related features**

Feature 5. Name: “sp_match_count”. Let $I(S_{i}\left( k \right)=S_{j}\left( k \right))$ be an indicator function, which equals 1 if the character in sequence *i*, column *k*, is a letter (not a gap character) and is identical to the letter in the same column in sequence *j*. The feature is then: $\sum_{i=1}^{N-1} \sum_{j=i+1}^{N} \sum_{k=1}^{L} I(S_{i}\left( k \right)=S_{j}\left( k \right))$.

Feature 6. Name: “sp_match_count_norm”. The feature is the same as feature 5, but normalized both for the number of pairs and the alignment length: $\frac{\sum_{i=1}^{N-1} \sum_{j=i+1}^{N} \sum_{k=1}^{L} I(S_{i}\left( k \right)=S_{j}\left( k \right))}{(\begin{matrix} N \\ 2 \end{matrix})L}$.

Feature 7. Name: “sp_mismatch_count”. Let $I_{2}\left( S_{i}\left( k \right)\neq S_{j}\left( k \right) \right)$ be an indicator function, which equals 1 if the characters in both sequences *i* and *j*, column *k*, are letters (both are not a gap character) and these letters are different. The feature is then: $\sum_{i=1}^{N-1} \sum_{j=i+1}^{N} \sum_{k=1}^{L} I_{2}\left( S_{i}\left( k \right)\neq S_{j}\left( k \right) \right)$.

Feature 8. Name: “sp_mismatch_count_norm”. The feature is the same as feature 7, but normalized both for the number of pairs and the alignment length: $\frac{\sum_{i=1}^{N-1} \sum_{j=i+1}^{N} \sum_{k=1}^{L} I_{2}\left( S_{i}\left( k \right)\neq S_{j}\left( k \right) \right)}{(\begin{matrix} N \\ 2 \end{matrix})L}$.

Feature 9. Name: “sp_go”. For each pair of sequences, we count the number of gap-opening events. For example, when aligning AKTTAC against -A--C-, there are three gap events (two of length 1, and one of length 2). So, in total for this pair there were three gap-opening events. Another example: A--C against A--G. In this case the pairwise alignment after removing columns in which the two sequences have gaps is AC against AG. The total number of gap-opening events is 0. Let $x_{ij}$ be the total number of gap-opening events between sequences *i* and *j*. The feature we compute is: $\sum_{i=1}^{N-1} \sum_{j=i+1}^{N} x_{ij}$.

Feature 10. Name: “sp_go_norm”. The normalized version of feature 9: $\frac{\sum_{i=1}^{N-1} \sum_{j=i+1}^{N} x_{ij}}{(\begin{matrix} N \\ 2 \end{matrix})L}$.

Feature 11. Name: “sp_ge”. For each pair of sequences, we count the number of gap-extension events. For example, when aligning AKTTAC against -A--C-, there are three gap opening events: two of length 1, and one of length 2. So, in total for this pair there were four gap-extension events (a total of four gap characters). Another example: A-LC against A--G. In this case the pairwise alignment after removing columns in which the two sequences have gaps is ALC against A-G. The total number of gap-extension events is 1. In fact, the total number of gap extension per pair of sequences, is the number of – characters after removing all gap-in-front-of-a-gap pairs. Let $x_{ij}$ be the total number of gap-extension events between sequences *i* and *j*. The feature we compute is: $\sum_{i=1}^{N-1} \sum_{j=i+1}^{N} x_{ij}$.

Feature 12: Name: “sp_ge_norm”. The normalized version of feature 11: $\frac{\sum_{i=1}^{N-1} \sum_{j=i+1}^{N} x_{ij}}{(\begin{matrix} N \\ 2 \end{matrix})L}$.

Feature 13. Name: “sp_match_BLOSUM62 ”. For each pair of sequences, the pairwise alignment score can be computed as match-score + mismatch-score + gap-opening score + gap extension score. The match score for a pairwise alignment is the sum over all cases in which the same amino acid is found in both sequences. The score of each match is taken from the BLOSUM62 matrix.

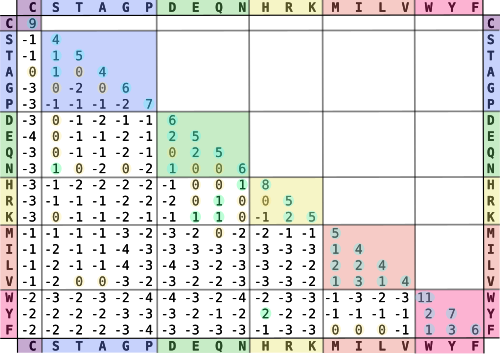

Thus, the match score for the following alignment, in which matches are in bold

**C**AM-**G**

**C**MEL**G**

is BLOSUM62(C,C)+ BLOSUM62(G,G) = 9+6 = 15.

The feature we compute is $\sum_{i=1}^{N-1} \sum_{j=i+1}^{N} x_{ij}$, where $x_{ij}$ is the pairwise match score between sequences *i* and *j*.

Feature 14. Name: “sp_match_norm_BLOSUM62”. Similar to feature 13, but normalized:
$\frac{\sum_{i=1}^{N-1} \sum_{j=i+1}^{N} x_{ij}}{(\begin{matrix} N \\ 2 \end{matrix})L}$ .

Feature 15. Name: “sp_mismatch_BLOSUM62”. This is very similar to the above “sp_BLOSUM62_match”, but here we sum over all mismatches in each pairwise alignment.

For the alignment, in which mismatches are in bold:

C**AM**-G

C**ME**LG

The pairwise mismatch score is BLOSUM62(A,M)+ BLOSUM62(M,E) = (-1)+(-2) = -3.

The feature we compute is $\sum_{i=1}^{N-1} \sum_{j=i+1}^{N} x_{ij}$ where $x_{ij}$ is the pairwise mismatch score between sequences *i* and *j*.

Feature 16. Name: “sp_mismatch_norm_BLOSUM62”. The normalized version of feature 15: $\frac{\sum_{i=1}^{N-1} \sum_{j=i+1}^{N} x_{ij}}{(\begin{matrix} N \\ 2 \end{matrix})L}$ where $x_{ij}$ is the pairwise mismatch score between sequences *i* and *j*.

Feature 17. Name: “sp_BLOSUM62_GO_-10_GE_-0.5”. This is the sum-of-pairs score assuming the BLOSUM62 with a gap opening score of 10 and a gap extension penalty of -0.5. The score is in fact: $\text{sp\_match\_}\text{BLOSUM62}+\text{sp\_mismatch\_}\text{BLOSUM62}-10\times"sp\_go"-0.5\times"sp\_ge"$.

Feature 18. Name: “sp_norm_BLOSUM62_GO_-10_GE_-0.5”. This is the same as feature 17, but normalized by the number of sequence pairs and length.

Feature 19. Name: “sp_norm_BLOSUM62_GO_-6_GE_-0.5”. This is the sum-of-pairs score assuming the BLOSUM62 with a gap opening score of 6 and a gap extension penalty of -0.5. The score is in fact: $\text{sp\_match\_}\text{BLOSUM62}+\text{sp\_mismatch\_}\text{BLOSUM62}-6\times"sp\_go"-0.5\times"sp\_ge"$.

Feature 20. Name: “sp_BLOSUM62_GO_-6_GE_-0.5”. This is the same as feature 19, but normalized by the number of sequence pairs and length.

Feature 21. Name: “sp_BLOSUM62_GO_-10_GE_-1”. This is the sum-of-pairs score assuming the BLOSUM62 with a gap opening score of 10 and a gap extension penalty of -1.0. The score is in fact: $\text{sp\_match\_}\text{BLOSUM62}+\text{sp\_mismatch\_}\text{BLOSUM62}-10\times"sp\_go"-1.0\times"sp\_ge"$.

Feature 22. Name: “sp_norm_BLOSUM62_GO_-10_GE_-1”. This is the same as feature 21, but normalized by the number of sequence pairs and length.

Feature 23. Name: “sp_BLOSUM62_GO_-6_GE_-1”. This is the sum-of-pairs score assuming the BLOSUM62 with a gap opening score of 6 and a gap extension penalty of --1. The score is in fact: $\text{sp\_}\text{BLOSUM62}\text{\_match}+\text{sp\_}\text{BLOSUM62}\text{\_mismatch}-6\times\text{sp\_go}-1.0\times"sp\_ge"$.

Feature 24. Name: “sp_norm_BLOSUM62_GO_-6_GE_-1”. This is the same as feature 23, but normalized by the number of sequence pairs and length.

Feature 25. Name: “sp_BLOSUM62_GO_-10_GE_-0.2”. Same as above, with a gap extension penalty of -0.2.

Feature 26. Name: “sp_norm_BLOSUM62_GO_-10_GE_-0.2”. This is the same as feature 26, but normalized by the number of sequence pairs and length.

Feature 27. Name: “sp_BLOSUM62_GO_-6_GE_-0.2”. Same as above, with a gap extension penalty of -0.2.

Feature 28. Name: “sp_norm_BLOSUM62_GO_-6_GE_-0.2”. This is the same as feature 27, but normalized by the number of sequence pairs and length.

Features 29-44. This is the same as features 13-28, but instead of using the BLOSUM62, we use the PAM250 matrix. The names of the features are from sp_PAM250_match to sp_PAM250_SOP_GO_6_GE_0.2_NORM.

Features 45-50. This is the same as features 17,19,21,23,25, and 27. The difference is that now, each pair of sequences has an associated weight. The sum-of-pairs here is $\sum_{i=1}^{N-1} \sum_{j=i+1}^{N} {w_{ij}x}_{ij}$ where $x_{ij}$ is the pairwise BLOSUM62 score (with the appropriate gap open and gap extension penalties) between sequences *i* and *j*. In this case $w_{ij}=w_{i}w_{j}$, where $w_{i}$ is the weight of sequence *i* according to the Henikoff weighting method (Henikoff and Henikoff 1994). Briefly, the method works as follows. For each column in the alignment, the number of different characters is computed. Here gap characters are treated as the 21th character (hence the “with_gaps” in the feature names). Assume, for example that among 20 sequences, 5 sequences have “M” in this position and 15 sequences have “K”. Because we have two characters, all the sequences with M divide a score of 0.5 between them and all sequences with K divide a score of 0.5 between them. Thus, each sequence with an M character gets a score of 0.5/5 and each sequence with a K character gets a score of 0.5/15. Now consider that the second position has 20 different characters. Each sequence gets a score of 1/20. This process of dividing the score is repeated for each position. The total score of a sequence is the sum of scores assigned for each from each position. This way, the more unique a sequence is, the higher its total score. The scores are normalized over all sequences so that they sum to 1.0. Name: from “sp_HENIKOFF_with_gaps_BLOSUM62_GO_-10_GE_-0.5” to “sp_HENIKOFF_with_gaps_BLOSUM62_GO_-6_GE_-0.2”.

Features 51-56. This is the same as features 45-50. The only difference is in the way the weights are computed. In this version, gap characters are ignored. Thus, if there is a column with 5 “M”’s, 12 “K”’s and 3 gap (“-“) characters, each M sequence gets 0.5/12 score, while if gaps were considered, each sequence harboring the M character would have received 0.333/5. The sequences with the gaps do not receive any score for that position. Name: “sp_HENIKOFF_without_gaps_BLOSUM62_GO_-10_GE_-0.5” to “sp_HENIKOFF_without_gaps_BLOSUM62_GO_-6_GE_-0.2”.

Features 57-68. This is the same as features 45-56. The only difference is that here, PAM250 is used instead of BLOSUM62. Name: “sp_HENIKOFF_with_gaps_PAM250_GO_-10_GE_-0.5” to “sp_HENIKOFF_without_gaps_PAM250_GO_-6_GE_-0.2”.

Features 69-74. This is the same as features 45-50. The only difference is that here, weights are computed based on the Clustal method (Julie D. Thompson et al. 1994). In the CLUSTAL weighting method, the weight of a leaf (i.e., a sequence) in a rooted tree is determined by aggregating the contributions from all branches along the path from the root to the leaf. Each branch contributes a value equal to its length divided by the number of leaf nodes (sequences) it subtends. Consequently, the computation of sequence weights necessitates a rooted tree. Because most trees obtained from tree search methods are unrooted, in this set of featuring, the midpoint rooting method is used, in which the root is placed at the midpoint of the longest path between any two leaves in the unrooted tree. In this set of features, the BLOSUM62 matrix is used. Names: “sp_CLUSTAL_WEIGHTS_mid_root_BLOSUM62_GO_-10_GE_-0.5” to “sp_CLUSTAL_WEIGHTS_mid_root_BLOSUM62_GO_-6_GE_-0.2”

Features 75-80. This is the same as features 69-74, but a different method is used to root the tree, the Differential Sum method (Julie D. Thompson et al. 1994). The first stage of this method involves computing, for each branch, the absolute difference (delta) between the total branch lengths on either side of that branch. A branch is considered a candidate for rooting if its own length exceeds this delta, indicating that it may lie on the optimal rooting path. In the second stage, among all candidate branches, the algorithm selects the one that minimizes the tree’s maximum depth — i.e., the branch that results in the shallowest tree, where the longest distance from the root to any leaf is minimized. Names from “sp_CLUSTAL_WEIGHTS_diff_sum_BLOSUM62_GO_-10_GE_-0.5 “ to “sp_CLUSTAL_WEIGHTS_diff_sum_BLOSUM62_GO_-6_GE_-0.2”.

Feature 81-92. This is similar to features 69-80, but here, the PAM matrix is used instead of BLOSUM. Name: “sp_CLUSTAL_WEIGHTS_mid_root_PAM250_GO_-10_GE_-0.5“ to “sp_CLUSTAL_WEIGHTS_diff_sum_PAM250_GO_-6_GE_-0.2”

**4. Gap related features**

Feature 93. Name: “num_gap_segments_norm”. A gap segment is one or more consecutive gap characters. For example, in the sequence L--C-T-----MMM there are 3 gap segments. This feature is the total number of gap segments divided by the number of sequences.

Feature 94. Name: “av_gap_segment_length”. This is the average length of a segment. It is the total number of gap characters in the MSA divided by the total number of gap segments.

Feature 95. Name: “gaps_len_one”. This is the total number of gap segments that are of length 1. The total is divided by *N,* the number of sequences in the MSA. In the following alignment:
C-M-G

C-MEL

CDM—

CDMQQ

The value is 0.75, as there are three segments of length 1, and *N* = 4.

Feature 96. Name: “gaps_len_two”. This is the total number of gap segments that are of length 2. The total is divided by *N,* the number of sequences in the MSA.

Feature 97. Name: “gaps_len_three”. This is the total number of gap segments that are of length 3. The total is divided by *N,* the number of sequences in the MSA.

Feature 98. Name: “gaps_len_four_plus”. This is the total number of gap segments that are of length 4 or more. The total is divided by *N,* the number of sequences in the MSA.

Feature 99. Name: “num_unique_gaps”. A unique segment is a gap segment that is unique to one sequence. Consider a gap that starts in alignment column i and end in alignment column j. It is considered unique if there is no other sequence that has a gap starting in column i and ending in column j. This feature counts the number of unique segments.

Feature 100. Name: “num_unique_gaps_norm”. This feature counts the number of unique segments, divided by *N,* the number of sequences in the MSA.

Feature 101. Name: “avg_unique_gap_length”. This feature is the average length of all unique segments across the MSA.

Feature 102. Name: “gaps_1seq_len1”. This feature is the number of gaps of length one, which are unique (appear in only a single sequence).

Feature 103. Name: “gaps_2seq_len1”. This feature is the number of gaps of length one, which appear in two sequences only.

Feature 104. Name: “gaps_1seq_len2”. This feature is the number of gaps of length two, which are unique.

Feature 105. Name: “gaps_2seq_len2”. This feature is the number of gaps of length two, which appear in two sequences only.

Feature 106. Name: “gaps_1seq_len3”. This feature is the number of gaps of length three, which are unique.

Feature 107. Name: “gaps_2seq_len3”. This feature is the number of gaps of length three, which appear in two sequences only.

Feature 108. Name: “gaps_1seq_len4plus”. This feature is the number of gaps of length four or more, which are unique.

Feature 109. Name: “gaps_2seq_len4plus”. This feature is the number of gaps of length four or more, which appear in two sequences only.

Feature 110. Name: “gaps_all_except_1_len1”. This feature is the number of gaps of length one, which are in all the sequences except one (they are missing in only a single sequence).

Feature 111. Name: “gaps_all_except_1_len2”. This feature is the number of gaps of length two, which are in all the sequences except one (they are missing in only a single sequence).

Feature 112. Name: “gaps_all_except_1_len3”. This feature is the number of gaps of length three, which are in all the sequences except one (they are missing in only a single sequence).

Feature 113. Name: “gaps_all_except_1_len4plus”. This feature is the number of gaps of length four or more, which are in all the sequences except one (they are missing in only a single sequence).

Feature 114. Name: “num_cols_no_gaps”. This feature is the number of MSA columns without a single gap character.

Feature 115. Name: “num_cols_1_gap”. This feature is the number of MSA columns with a single gap character. Consider the following MSA:

C-M-G

C-MEL

CDM—

CDMQQ

This feature equals 1 because only the last column has a single gap character.

Feature 116. Name: “num_cols_2_gaps”. This feature is the number of MSA columns with exactly two gap characters.

Feature 117. Name: “num_cols_all_gaps_except1”. This feature is the number of MSA columns in which all the characters are gaps, except one.

Feature 118. Name: “single_char_count”. This feature is the number of cases of a single character that has either a gap or a sequence start /end in both sides of it. In other words, we go over each sequence and count the number of cases in which we see “-C-“, in which C represent a non-gap character. We also count cases such as “C-M…”, that is, the sequence starts with a non-gap character followed by a gap. Finally, we also count cases such as “MK-Q “, in which the sequence ends with a a single gap character followed by a non-gap character at the end of the sequence. Consider the following MSA:

**C-M-G**

**C-**MEL

CDM—

CDM**-Q**

This feature equals 4 because in the first sequence we have “C-“ in the beginning of the sequence, “-M-“ in the middle of the sequence, and “-G“ at the end; in the second sequence we have “C-“, and in the last sequence we have a “-Q” at the end.

Feature 119. Name: “double_char_count”. This feature is the number of cases of exactly two characters that have either a gap in both sides of a gap in one side and a sequence start or end at the other side. The sequence “AA-CTT---MM--QQQ-LL” has three such cases (AA-, -MM- and -LL”.

Feature 120. Name: “n_unique_sites”. Some MSA columns are identical. This feature counts the number of columns in the MSA, after filtering identical columns.

**5. Tree related features**

From the given MSA, a NJ tree is built. The pairwise distances are computed using Kimura’s distance for proteins: $d=-ln(1-p-0.2p^{2})$, where $p$ is the fraction of mismatches between the two sequences (Kimura 1983). Note that positions with gaps are considered when computing $p$. A NJ tree is built from these distances (using an in-house python script). The result is an unrooted tree with *N* sequences and branches $\vec{b}=(b_{1},\ldots,b_{2N-3})$.

Feature 121. Name: “bl_mean”. The mean of the vector $\vec{b}$. The prefix, “bl”, stands for branch lengths.

Feature 122. Name: “bl_sum”. The sum of the elements in vector $\vec{b}$.

Feature 123. Name: “bl_max”. The length of the longest branch (the max over the entries in vector $\vec{b}$).

Feature 124. Name: “bl_min”. The length of the shortest branch (the min over the entries in vector $\vec{b}$).

Feature 125. Name: “bl_25_pct”. The 25% percentile of the entries in vector $\vec{b}$.

Given a vector $\vec{b}$, the q^th^ percentile of $\vec{b}$ is the value q/100 of the way from the minimum to the maximum in a sorted copy of $\vec{b}$. The values and distances of the two nearest neighbours will determine the percentile if the normalized ranking does not match the location of q exactly.

when the desired percentile lies between two indexes $i$ and $j = i + 1$, we first determine $i + g$, a virtual index that lies between $i$ and $j$, where $i$ is the floor and $g$ is the
fractional part of the index. The final result is, then, an interpolation of $\vec{b}_{i}$ and $\vec{b}_{j}$ based on $g$. During the computation of $g$, $i$ and $j$, are modified using correction constants $\alpha$ and
$\beta$, according to the formula: $i + g = (q / 100) * ( n - \alpha- \beta+ 1 ) + \alpha$.

We use $\alpha=\beta=1$ according to method 7 in Hyndman and Fan (Hyndman and Fan 1996).

For example, for $\vec{b}=[23, 5, 3, 6, 9, 14, 1, 43]$, after sorting in an ascending order the vector is $\vec{b}=[1, 3, 5, 6, 9, 14, 23, 43]$. The 25th percentile value is 4.5.

Feature 126. Name: “bl_75_pct”. The 75% percentile of the entries in vector $\vec{b}$.

Based on the reconstructed NJ tree, the parsimony score is computed for each alignment column using the Fitch algorithm. The result is a vector $\vec{p}=(p_{1},\ldots,p_{L})$, where *L* is the number of alignment columns. We expect that the minimum entry for this vector would be zero, representing columns without substitutions. This feature is already accounted for.

Feature 127. Name: “parsimony_mean”. The mean of the vector $\vec{p}$.

Feature 128. Name: “parsimony_sum”. The sum of the elements in vector $\vec{p}$.

Feature 129. Name: “parsimony_max”. The max over the entries in vector $\vec{p}$.

Feature 130. Name: “parsimony_min”. The min over the entries in vector $\vec{p}$ (i.e., number of columns with a single type of letters, with or without gaps).

Feature 131. Name: “parsimony_25_pct”. The 25% percentile of the entries in vector $\vec{p}$.

Feature 132. Name: “parsimony_75_pct”. The 75% percentile of the entries in vector $\vec{p}$.

**6. Entropy related features**

The entropy of a discrete distribution is defined as $H\left( x \right)=-\sum_{x} p(x)\log_{2} p(x)$. The contribution of events with probability 0 is defined to be 0. For a column, $p(M)$ is the frequency of amino acid *M*. Of note, we do not assume pseudo-counts when computing the frequencies and we do not consider gaps as characters. We compute the entropy of each column, resulting in a vector $\vec{h}=(h_{1},\ldots,h_{L})$, where *L* is the number of alignment columns. Note that columns with parsimony score of zero will also have entropy of zero, so this is why the minimum over this $\vec{h}$ vector is not considered as a feature.

Feature 133. Name: “entropy_mean”. The mean of the vector $\vec{h}$.

Feature 134. Name: “entropy_sum”. The sum of the elements in vector $\vec{h}$. This is also called ES in the manuscript.

Feature 135. Name: “entropy_max”. The max over the entries in vector $\vec{h}$.

Feature 136. Name: “entropy_25_pct”. The 25% percentile of the entries in vector $\vec{p}$.

Feature 137. Name: “entropy_75_pct”. The 75% percentile of the entries in vector $\vec{p}$.

Feature 138. Name: “constant_sites_pct”. A column is considered as a “constant_sites “column if all the characters along the entire column are identical, where gaps are not considered as characters. This column can be also defined as a zero-entropy column. This feature calculates the percentage of “constant_sites “column out of the total number of columns. This is also called “column score” (CS) in the manuscript.

**7. kmer related features**

Consider all kmer of length *K*. We first generate a histogram by counting the occurrences of all substrings of length K, including gaps. The resulting output is a list of k-mers that are present in the MSA and their frequency. Consider the following MSA with *K*=4:

C-M-G

- CMEL

CC-M-

The K-mers are C-M- (frequency 2), -M-G (frequency 1), -CME (frequency 1), CMEL (frequency 1), and CC-M (frequency 1). The frequencies can be arranged in a vector $\vec{m}=(h_{1},\ldots,h_{N})$. In the above example the vector would be: (2,1,1,1,1).

Feature 139. Name: “k_mer_average_K5”. We generate the $\vec{m}$ vector with *K*=5 and take the average frequency. If the frequencies are (2,1,1,1,1), the average would be 6/5 = 1.2

Feature 140. Name: “k_mer_max_K5”. We generate the $\vec{m}$ vector with *K*=5 and take the maximal frequency.

Feature 141. Name: “k_mer_90_pct_K5”. We generate the $\vec{m}$ vector with *K*=5 and take the 90% percentile of the entries in vector $\vec{m}$.

Feature 142. Name: “k_mer_95_pct_K5”. We generate the $\vec{m}$ vector with *K*=5 and take the 95% percentile of the entries in vector $\vec{m}$.

Feature 143. Name: “kmer_sum_of_top_10_K5”. We generate the $\vec{m}$ vector with *K*=5 and take the sum of the 10 highest entries in vector $\vec{m}$.

Feature 144. Name: “k_mer_average_K10”. We generate the $\vec{m}$ vector with *K*=10 and take the average frequency.

Feature 145. Name: “k_mer_max_K10”. We generate the $\vec{m}$ vector with *K*=10 and take the maximal frequency.

Feature 146. Name: “k_mer_90_pct_K10”. We generate the $\vec{m}$ vector with *K*=10 and take the 90% percentile of the entries in vector $\vec{m}$.

Feature 147. Name: “k_mer_95_pct_K10”. We generate the $\vec{m}$ vector with *K*=10 and take the 95% percentile of the entries in vector $\vec{m}$.

Feature 148. Name: “kmer_sum_of_top_10_K10”. We generate the $\vec{m}$ vector with *K*=10 and take the sum of the 10 highest entries in vector $\vec{m}$.

Feature 149. Name: “k_mer_average_K20”. We generate the $\vec{m}$ vector with *K*=20 and take the average frequency.

Feature 150. Name: “k_mer_max_K20”. We generate the $\vec{m}$ vector with *K*=20 and take the maximal frequency.

Feature 151. Name: “k_mer_90_pct_K10”. We generate the $\vec{m}$ vector with *K*=20 and take the 90% percentile of the entries in vector $\vec{m}$.

Feature 152. Name: “k_mer_95_pct_K20”. We generate the $\vec{m}$ vector with *K*=20 and take the 95% percentile of the entries in vector $\vec{m}$.

Feature 153. Name: “kmer_sum_of_top_10_K20”. We generate the $\vec{m}$ vector with *K*=20 and take the sum of the 10 highest entries in vector $\vec{m}$.

**8. Structure-related features**

To get the structure-related features, we first downloaded the ProstT5 model and weights using the Foldseek Linux AVX2 build (van Kempen et al. 2024), with the following command line:

*foldseek databases ProstT5 weights </path/to/prostt5/weights>*

Given an MSA, we then create a ProstT5-encoded database from its unaligned sequences FASTA file with the command:

*foldseek createdb <unaligned_fasta_file> <embeddings_database_dir> --prostt5-model* < */path/to/prostt5/weights* > *--threads 128.*

This database is a set of protein sequences that have been transferred into high-dimensional vector representations, known as ProstT5 embeddings, using the Protein structure-sequence T5 (ProstT5) language model. These embeddings capture structural and evolutionary features and are stored in Foldseek’s optimized, indexed database format to enable fast similarity searches and alignments. The ProstT5 encoder is used instead of its default 3Di structural encoding because the input is not structural, but rather amino-acid sequences.

We then used FoldMason Linux release 2-7bd21ed (Gilchrist et al. 2024), which utilizes the ProstT5-encoded Foldseek database to construct a structural MSA of our sequences:

*foldmason structuremsa {embeddings_database_dir} {output_structural_MSA_dir}*

Now we have two MSAs, the original MSA for which features are being extracted, and the structural MSA generated by FoldMason. Two features were then calculated from these two MSAs: (1) a fraction of columns in the reference alignment that is present in the structural alignment (column score); and (2) similarly, the fraction of residue pairs that are aligned together in both alignments (residue pairs score).
